# Integrating disease and protein networks expands genetic support for therapeutic targets

**DOI:** 10.64898/2026.11.17.767419

**Authors:** Wanheng Zhang, Jie Xu, Yue Lyu, Minghui Liu, Hui Shen, Lang Wu, Bingxin Zhao, Hong-Wen Deng, Chong Wu

## Abstract

Human genetics guides therapeutic target selection, but direct genetic support covers few clinically investigated target–indication pairs, each linking a target to a disease. Here we introduce BRIDGE (Biological Relation–Informed Diffusion of Genetic Evidence), which propagates genetic evidence through coupled protein–protein interaction and disease-similarity networks. Each candidate’s score is calibrated against scores obtained after randomly substituting similarly connected genes for genes with direct genetic support. In a retrospective benchmark of 13,022 pairs that had reached at least phase I, BRIDGE supported 572 additional pairs, increasing coverage from 5.1% to 9.5%. Among eligible pairs lacking direct genetic support, 25.3% of network-added pairs had reached launch versus 8.6% of remaining candidates (relative success, 2.94; 95% confidence interval, 2.51–3.45). This enrichment extended to 158 pairs involving 101 genes with no direct genetic support in the benchmark. At a matched selection size of 1,310 pairs, BRIDGE ranking captured 265 launched pairs, compared with 223 under a rule transferring support across neighbouring targets or indications. These findings show that coupling disease relationships with protein connectivity expands the reach of human genetics to additional therapeutic targets and indications while retaining clinical relevance.

## Introduction

Human genetics provides a basis for therapeutic target selection, with supported mechanisms more likely to progress through clinical development.^1–5^ Genetics-guided scores track clinical stage,^6^ and trials lacking strong genetic support more often stop for insufficient efficacy.^7^ Yet even with systematic mapping of associated genetic regions to genes (locus-to-gene mapping),^8^ direct genetic support requires genetic evidence linking a specific gene to a trait that closely matches the intended treatment indication.^5^ A target–indication (T–I) pair combines a therapeutic target, represented here by its gene, with a disease or condition to be treated. Many such pairs under clinical development therefore lack direct genetic support. Extending this evidence to interacting targets or related indications could broaden the utility of human genetics. The central challenge therefore is to expand its reach without diluting its clinical relevance.

Protein–protein interaction (PPI) and disease-similarity networks offer complementary routes for this extension. Protein connectivity can link a candidate target to genes with direct genetic support for the same indication, whereas disease relationships can provide access to evidence from related indications. Network propagation spreads scores along network connections to prioritize genes. Random-walk methods combine different types of connections to recover disease genes whose known associations were hidden during evaluation.^9–13^ Analyses combining genetics with protein networks have also identified therapeutic targets and linked genetic associations to groups of functionally related proteins.^14–16^ These approaches provide a basis for extending genetic evidence across both targets and indications.

The clinical value of this expansion, however, cannot be inferred from the recovery of known disease genes alone. Clinical enrichment means that supported pairs have a higher proportion of successful development outcomes than comparison pairs. Simple PPI-neighbour and pathway expansions have shown weaker clinical enrichment than targets with direct genetic support, whereas protein-complex expansion retained enrichment with smaller gains in coverage.^15^ Relaxing trait–indication matching can also weaken the association with clinical success.^5^ Biases favouring highly connected genes^17^ and precision–sensitivity trade-offs^18^ further motivate controls that account for gene connectivity and make propagation scores comparable.^19,20^ Whether coupling protein and disease networks can expand genetic support while preserving clinical enrichment across diverse targets and indications remains unknown. A comprehensive assessment should therefore establish both the clinical relevance of the added candidates and whether integrating these relationships improves prioritization beyond their connections to existing genetic evidence.

To address these challenges, here we introduce BRIDGE (Biological Relation–Informed Diffusion of Genetic Evidence), which propagates genetic evidence through coupled PPI and disease-similarity networks. Each candidate’s score is compared with scores obtained after substituting similarly connected genes for genes with direct genetic support to assign network support. In a retrospective benchmark of 13,022 T–I pairs that had reached at least phase I, combining direct genetic support and network-added support increased coverage from 5.1% to 9.5%, extending support to genes outside the direct seed set, while retaining enrichment for clinical success. BRIDGE captured more launched pairs than the local-transfer rule at matched selection size. Seed-withholding analyses revealed complementary contributions of disease relationships and protein connectivity to evidence transfer. Finally, therapeutic case studies linked network-added candidates to clinically investigated pathways. Together, these analyses show how integrating disease context and protein connectivity can broaden clinically informative genetic support for therapeutic target prioritization.

## Results

### BRIDGE expands genetic support while retaining clinical enrichment

BRIDGE extends genetic support through network propagation followed by pair-specific empirical calibration (**Fig. 1a**). Its two-layer network contains gene nodes and indication nodes. Gene nodes represent their encoded proteins and are linked in a PPI network. Its edges are high-confidence functional associations from STRING v12.0, a database of protein associations (combined confidence ≥0.90)^21^. The disease layer links indications using Medical Subject Headings (MeSH), a standardized biomedical vocabulary, with similarity ≥0.50. T–I pairs with direct genetic support from the Minikel *et al.* benchmark^5^ provide genetic seed assignments and gene–disease links connecting the two layers. Genes in these assignments are seed genes; the same gene can be a seed for one indication and a candidate for another.

**Figure 1.**
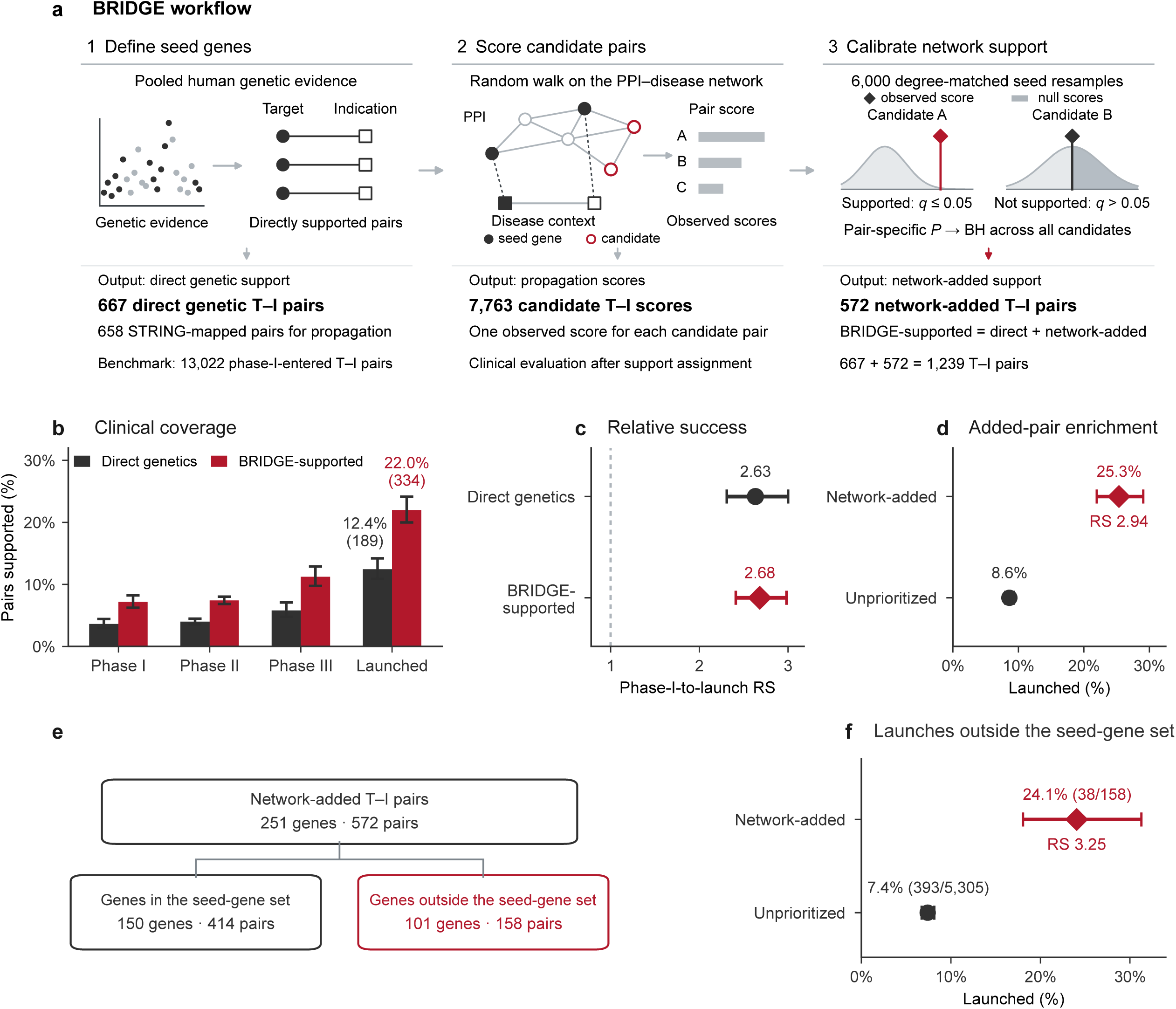
BRIDGE expands genetic support while retaining clinical enrichment. **a,** Workflow for coupled protein–protein interaction (PPI) and disease-network propagation, followed by pair-specific empirical calibration. BRIDGE-supported pairs (the full BRIDGE support set) combine direct genetic support and network-added support; the evidence, network and score-distribution graphics are schematic. **b,** Supported proportions within mutually exclusive clinical-status strata: phase I (*n* = 2,569), phase II (*n* = 7,383), phase III (*n* = 1,551) and launched (*n* = 1,519). Phase II and III combine resolved and active pairs. Black and crimson bars denote direct genetic support and full BRIDGE. **c,** Phase-I-to-launch relative success (RS; ratio of launch proportions) for direct genetic support (*n* = 667) and full BRIDGE (*n* = 1,239), each compared with its complement in the 13,022-pair cohort. Points and lines show RS and Katz confidence intervals (CIs); the dotted line marks RS = 1. **d,** Launch proportions for network-added (*n* = 572; crimson diamond) and unprioritized (*n* = 7,191; black circle) candidates. **e,** Network-added genes and pairs partitioned by membership in the 359-gene mapped seed set. Labels give gene and pair counts. **f,** Launch proportions for candidates whose genes lie outside the seed set: network-added (*n* = 158) and unprioritized (*n* = 5,305), with the same symbols as d. Lines in b, d and f show Wilson CIs; annotations in d and f give RS. All CIs are two-sided 95% intervals. Calibration and comparator definitions are given in Methods. Source data are provided as a Source Data file.

For each focal indication, the indication currently being evaluated, a random walk with restart^13^ begins at its disease node, follows network connections and returns to the starting node with a fixed probability. The resulting propagation score measures each candidate target’s network connection to that indication. Eligible candidates are benchmark pairs that map to the network, belong to an indication containing genetic seeds, and lack direct genetic support for that indication.

We compare each candidate’s observed score with its own null distribution, a random reference generated from 6,000 degree-matched seed resamples. Resampling replaces seed genes with genes of similar weighted PPI degree, the sum of a gene’s PPI edge weights, while keeping PPI and disease-similarity edges fixed. One-sided empirical values quantify how often resampling yields a score at least as high as observed. Joint adjustment across all eligible non-seed pairs produces q values using the Benjamini–Hochberg (BH) procedure^22^, with q ≤ 0.05 defining network-added support (**Methods**).

We retrospectively evaluated BRIDGE in the Minikel *et al.* benchmark^5^ of 13,022 T–I pairs that had reached at least phase I. The 1,519 launched pairs had a drug reach launch for the corresponding target and indication. 667 pairs had direct genetic support; 658 mapped to the PPI network and served as seeds across 241 indications. Together, these seed pairs defined the seed-gene set: 359 genes with direct genetic support for at least one indication. Among 7,763 eligible non-seed candidates, BRIDGE designated 572 pairs as network-added; the remaining 7,191 candidates were unprioritized. Combining direct genetic support and network-added support produced full BRIDGE, the union of these support sets (1,239 BRIDGE-supported pairs). Coverage, the fraction of all 13,022 benchmark pairs receiving support, increased from 5.1% to 9.5%. Full BRIDGE supported 334 T–I pairs for which a drug had reached launch, compared with 189 with direct genetic support (77% increase; **Fig. 1b**).

The expanded set retained clinical enrichment. Relative success (RS) divides the proportion reaching a specified clinical outcome in the supported group by that in the comparison group. Values above 1 indicate enrichment. Full BRIDGE had a phase-I-to-launch RS of 2.68 (95% confidence interval (CI), 2.41–2.98), compared with 2.63 (95% CI, 2.31–3.00) for pairs with direct genetic support (**Fig. 1c**). Each comparison used the remaining pairs in the full benchmark as its reference group. Network-added pairs were also clinically enriched, with an RS of 2.94 (95% CI, 2.51–3.45). Among these pairs, 145 of 572 had reached launch (25.3%), compared with 620 of 7,191 unprioritized non-seed candidates (8.6%; **Fig. 1d** and **Supplementary Table 2**).

BRIDGE also expanded support to genes outside the mapped direct genetic seed set. Among the 251 genes represented in the network-added pairs, 101 had no direct genetic seed assignment in any indication. Adding these 101 genes increased the mapped supported gene set from 359 to 460 (28.1% expansion; **Fig. 1e**). Among candidates whose genes lay outside the seed set, 38 of 158 supported pairs had launched, compared with 393 of 5,305 unprioritized pairs (24.1% versus 7.4%; RS, 3.25; 95% CI, 2.42–4.35; **Fig. 1f**, **Supplementary Table 3** and **Methods**).

Compared with direct genetic support, full BRIDGE supported more launched T–I pairs across all 17 therapy areas (**Supplementary Fig. 1a**). Despite expanding coverage, BRIDGE generally preserved the phase-I-to-launch enrichment observed for direct genetic support across therapy areas (**Supplementary Fig. 1b**). Clinical enrichment also persisted across the evaluated development transitions, including phase I→II, phase II→III, phase III→launch, and overall phase I→launch progression (**Supplementary Fig. 1c**). Enrichment also persisted among targets launched or under development in other indications (**Supplementary Fig. 9**). Finally, the pattern was robust across six genetic-evidence sources: Online Mendelian Inheritance in Man (OMIM), Genebass, Open Targets Genetics genome-wide association study (GWAS) evidence, augmented Open Targets genetics, PICCOLO expression quantitative trait locus (eQTL) evidence and pooled genetics (**Supplementary Fig. 1d** and **Supplementary Table 2**).

Across sensitivity analyses varying the STRING confidence threshold, disease-similarity threshold, restart probability, the weight of transitions between layers (inter-layer coupling), and the support threshold, full BRIDGE continued to expand coverage while retaining clinical enrichment (**Supplementary Fig. 2)**. Similar patterns were observed after removing STRING evidence extracted from scientific text (the text-mining channel) and recalibrating the resulting PPI network (**Supplementary Fig. 2**, **Supplementary Table 2** and **Methods**).

### Disease context and protein connectivity extend genetic evidence across indications and targets

We examined how PPI connectivity and disease relationships contributed to BRIDGE’s expanded genetic support. We compared three propagation architectures using the same genetic seed assignments, with separate empirical calibration for each architecture (**Methods**). PPI-only propagation, which spread evidence through protein interactions from seeds for the focal indication, supported 52 pairs (RS, 1.96). Adding gene–disease links connecting genes to the indications, which allows the random walk passes from a gene to one of its genetically supported indications and backs to other seed genes for that indication. This expansion increased support to 209 pairs (RS, 3.47). Further adding disease–disease similarity links to form the full BRIDGE network expanded support a further 2.7-fold to 572 pairs, an 11-fold expansion over PPI-only propagation (**Fig. 2a**). The expanded set remained enriched for launches (RS, 2.94; **Fig. 2b**).

**Figure 2.**
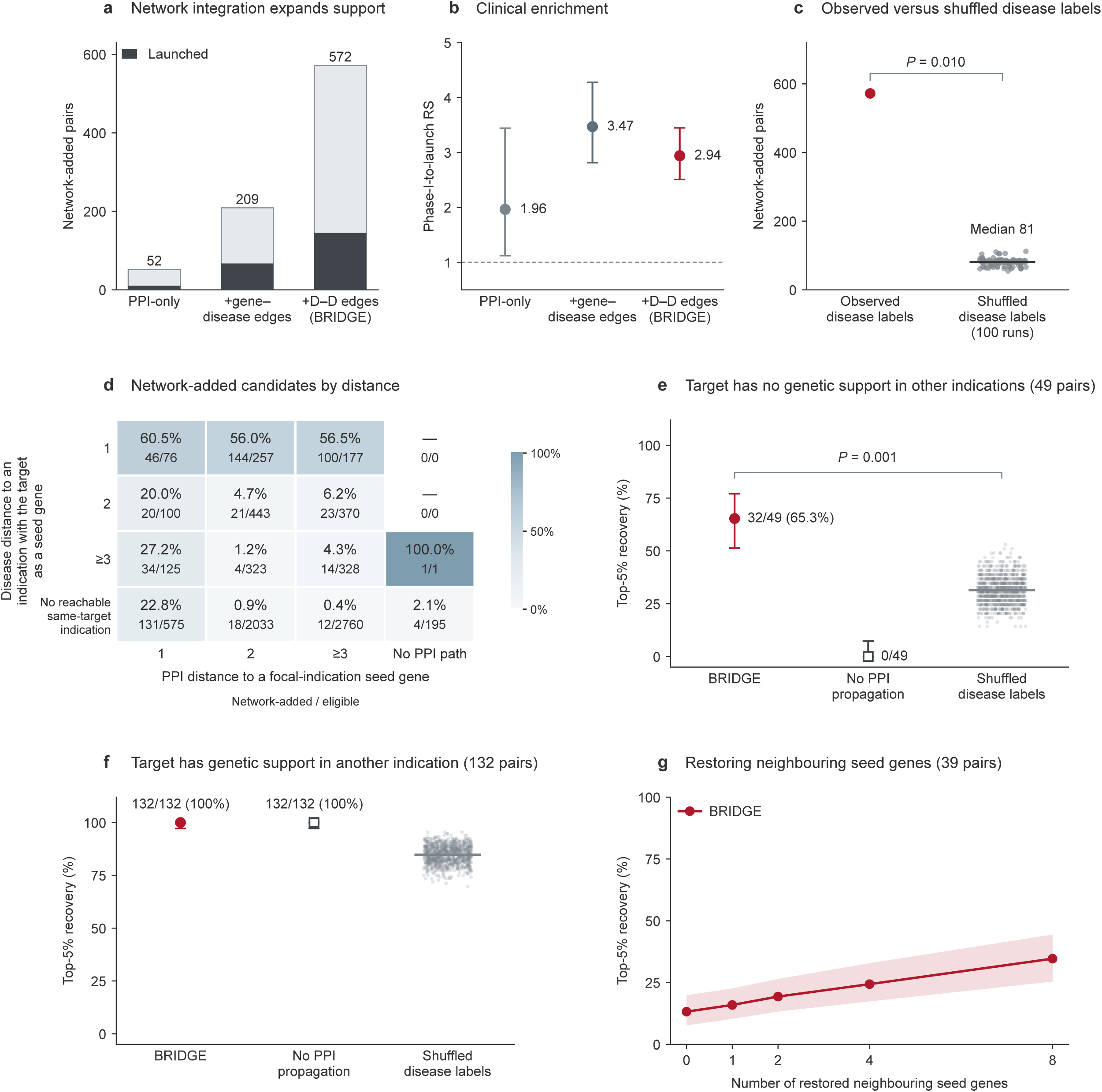
Disease context and protein–protein interaction (PPI) connectivity enable complementary evidence transfer. **a,** Calibrated support among 7,763 non-seed pairs across three propagation architectures; D–D denotes disease–disease edges. Each used the same 658 seed assignments, calibrated separately at Benjamini–Hochberg *q* ≤ 0.05 (Methods). Dark grey, launched pairs; pale grey, other supported pairs. **b,** Relative success (RS), comparing launch proportions in supported pairs with each architecture’s remaining candidates. Points and lines show RS and Katz log-scale confidence intervals (CIs). **c,** Observed support versus 100 recalibrated disease-label shuffles; the horizontal line marks the shuffle median. **d,** BRIDGE support by disease and PPI distance. Disease distance is the shortest path from the focal indication to an indication seeding the same target, using disease edges with Medical Subject Headings (MeSH) similarity ≥0.50. PPI distance counts edges in the shortest path from the target to a focal-indication seed gene. Each axis shows distances of 1, 2 and ≥3 steps or no path (“No reachable same-target indication”; “No PPI path”). Colours and percentages show supported proportions; counts give supported/eligible pairs; a dash (0/0) marks cells without eligible pairs. **e,f,** Target recovery after withholding all focal-indication seeds, for launched pairs whose targets are unseeded elsewhere (**e**, *n* = 49) or remain seeded elsewhere (**f**, *n* = 132). The No PPI propagation control replaces the PPI transition matrix with the identity while retaining disease and gene–disease transitions. Points and lines show recovery and Wilson CIs; grey dots show 1,000 disease-label permutations (horizontal lines, means). Brackets in c and e report one-sided empirical *P* values. **g,** BRIDGE recovery after restoring 0, 1, 2, 4 or 8 original seed genes from directly neighbouring indications (39 pairs across 33 indications with at least eight available anchors). Focal-indication seeds remain withheld; zero anchors denotes the degree-matched replacement background (Methods). Points average 50 backgrounds within each pair, then pairs; shading shows pointwise percentile CIs from 5,000 indication-cluster bootstrap resamples. Recovery in e–g uses the top 5% of 19,699 genes. All CIs are two-sided 95% intervals; Wilson intervals treat pairs as independent. Source data are provided as a Source Data file.

Adding network links creates more routes for propagation, so we tested whether the expansion depended on the specific disease relationships. We shuffled disease labels while preserving the disease network’s structure. Across 100 independently recalibrated shuffles, support fell to 55–112 pairs (median, 81; empirical *P* = 0.010; **Fig. 2c**), indicating that the alignment between disease relationships and genetic evidence contributed to the expansion. The 248 observed network-added pairs not recovered in any shuffle also retained launch enrichment (50 launched pairs, 20.2%; **Supplementary Fig. 3**).

We next asked how BRIDGE support related to the connections between individual candidates and existing genetic evidence. To characterize these routes, we stratified all 7,763 non-seed candidate T–I pairs by their proximity to genetic evidence along both dimensions (**Fig. 2d**). Distance was the number of edges in the shortest path, with each edge counted as one step. PPI distance linked each target to the nearest genetic seed for the focal indication; disease-network distance linked the focal indication to the nearest indication with direct genetic support for the same target.

Support patterns revealed two complementary routes of evidence transfer. First, BRIDGE supported same-target transfer, defined here as direct genetic support for the same target in a directly neighbouring indication, one disease-similarity edge away. Among candidate T–I pairs eligible for same-target transfer (*n* = 510), BRIDGE support rates ranged from 56.0% to 60.5% across different PPI distances to genetic seeds for the focal indication, including targets at least three PPI steps from a focal-indication seed (**Fig. 2d**). This pattern indicates that same-target transfer can support candidates even when the target is molecularly distant from genetic seeds for the indication.

Second, among the remaining 7,253 candidate T–I pairs, direct genetic support for the same target was at least two disease steps away or absent from all indications reachable from the focal indication through a disease-network path. In this group, BRIDGE supported 20.0–27.2% of candidates with a direct PPI connection to a genetic seed for the focal indication, compared with 0.4–6.2% of candidates two or more PPI steps away. BRIDGE also supported 5 of 196 candidates without a PPI-only path to a focal-indication seed (**Fig. 2d**), demonstrating that support could extend through connections unavailable within the PPI layer alone.

The typical genetic evidence support analysis required genetic evidence for the focal indication. We thus asked whether BRIDGE could recover launched targets for the focal indication only using genetic evidence assigned to other indications by withholding all genetic seed assignments for the focal indication. Eligibility required at least one directly neighbouring indication with remaining genetic seeds. Here, recovery meant ranking the withheld target within the top 5% of all 19,699 mapped genes (**Methods**).

We first examined 49 launched T–I pairs whose targets were not genetic seeds in any other indication. Recovery of these targets therefore required BRIDGE to access evidence from a related indication and then transfer it across targets through the PPI network. Under this strict setting, BRIDGE recovered 32 pairs (65.3%), whereas disabling PPI propagation reduced recovery to zero (**Fig. 2e**). Moreover, BRIDGE’s recovery rate was higher than any of the 1,000 runs with shuffled disease labels (mean, 31.4%; maximum, 53.1%; empirical *P* = 0.001). By contrast, when targets remained genetic seeds in other indications (132 launched T–I pairs), both BRIDGE and the version without PPI propagation recovered all 132 pairs (**Fig. 2f**).

We next tested whether cross-target recovery improved as more anchors became available. In this experiment, real anchors were genes other than the target that were genetic seeds for an indication directly neighbouring the focal indication. For example, when target X was tested for indication A, gene Y, which has direct genetic support for neighbouring indication B, served as an anchor. We first replaced every anchor with a random gene matched for weighted PPI degree, so the number of seeds and their degree distribution stayed the same. With focal-indication seeds still withheld, we then restored 0, 1, 2, 4 or 8 randomly selected real anchors and repeated this across 50 random replacement backgrounds (Methods). Mean top-5% BRIDGE recovery increased from 13.2% with no real anchors to 34.7% with eight (**Fig. 2g**; **Supplementary Fig. 4b–4d**). To test whether additional anchors helped when the nearest anchor was already available, we also retained that anchor throughout restoration. In this setting, recovery increased by 6.3 percentage points from one to eight anchors (**Supplementary Fig. 4c**).

Together, these analyses show that BRIDGE expands genetic support by combining evidence transfer across indications with propagation between targets. Disease relationships provide access to genetic evidence from other indications, while PPI propagation extends that evidence to additional targets, including those with no remaining seed assignments.

### BRIDGE refines target–indication prioritization beyond local-transfer rules

The preceding analyses showed that BRIDGE support often lay close to existing genetic evidence. Specifically, 83.0% of network-added pairs (475/572) had direct genetic support for the same target in a neighbouring indication or a direct PPI connection to a focal-indication seed (**Fig. 2d**). We therefore asked whether BRIDGE added useful discrimination beyond selecting candidates with these local connections. We defined a local-transfer rule among the same 7,763 non-seed candidates. A candidate was selected if either (i) the same target had direct genetic support in a neighbouring indication (same-target transfer; MeSH similarity ≥ 0.50) or (ii) the candidate target was connected by one PPI edge to a genetic seed for the same indication.

The local-transfer rule selected 1,310 pairs, including 475 of the 572 network-added pairs (**Fig. 3a**). Among the pairs selected by the local-transfer rule, launch proportions were 27.2% for network-added pairs and 11.3% for the remaining pairs (RS, 2.41; 95% CI, 1.82–3.24). Among the 6,453 pairs not selected by the rule, launch proportions were also higher for the 97 network-added pairs than for the remaining candidates (16.5% versus 8.3%; RS, 1.99; 95% CI, 1.12–3.02; **Fig. 3b**). Thus, BRIDGE identified clinically enriched subsets both within and beyond candidates selected by these local relationships.

**Figure 3.**
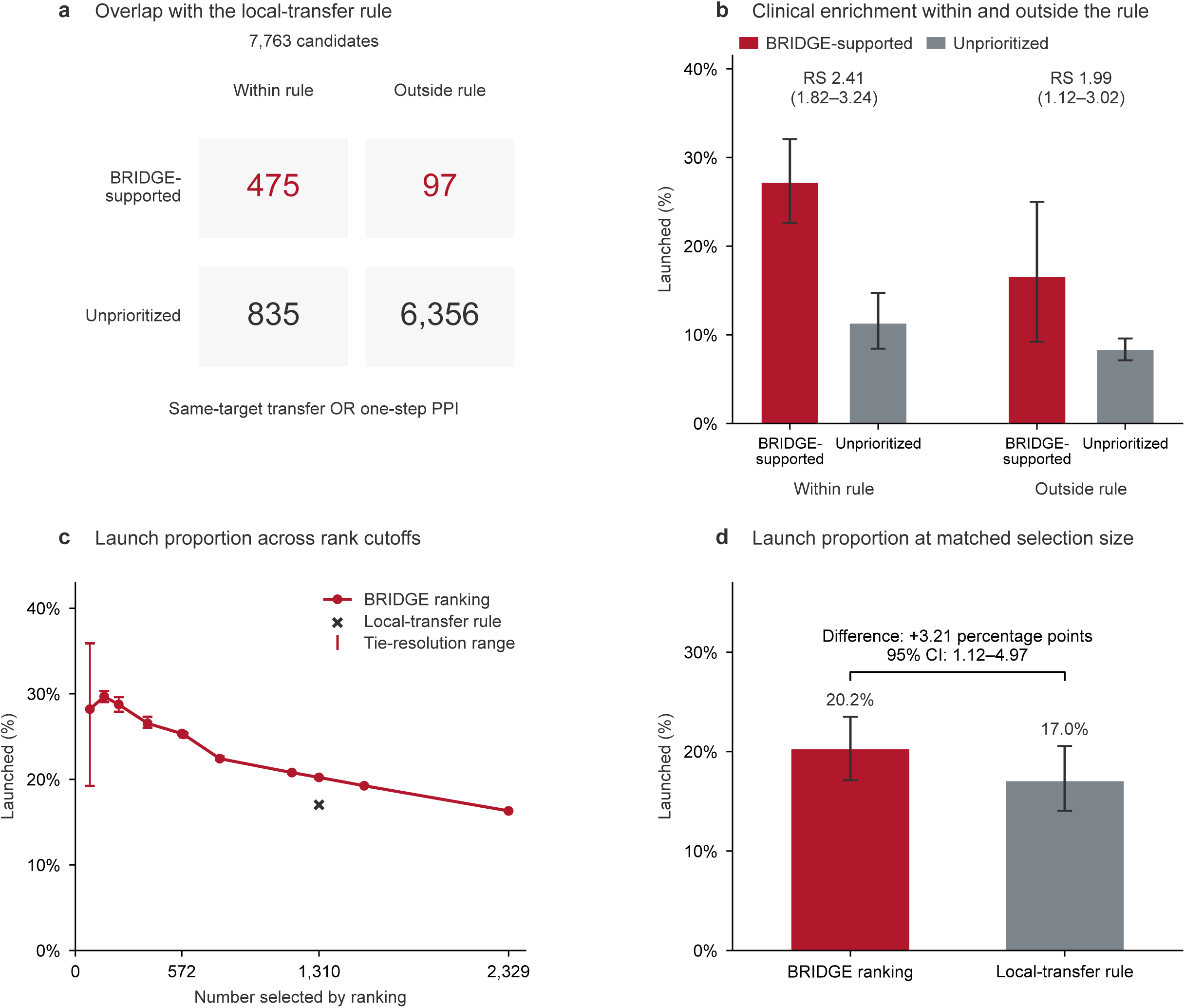
Target–indication prioritization beyond local-transfer rules. **a,** Membership of 7,763 non-seed pairs in BRIDGE’s calibrated support set and the local-transfer rule. The rule requires direct genetic support for the same target in a neighbouring indication (Medical Subject Headings (MeSH) similarity ≥0.50) or one protein–protein interaction (PPI) edge to a seed gene for the focal indication. **b,** Launch proportions among network-added (labelled BRIDGE-supported in the panel; burgundy) and unprioritized (grey) pairs within and outside the rule. Group sizes appear in a; launched counts, in bar order, are 129, 94, 16 and 526. Whiskers show 95% indication-cluster bootstrap confidence intervals (CIs). Relative success (RS) compares supported with unprioritized pairs within each stratum; parentheses give 95% CIs. **c,** Launch proportions across 11 selection depths for BRIDGE’s empirical-*P* ranking. The black cross marks the local-transfer rule’s operating point (1,310 pairs). The curve uses identifier-resolved ties; burgundy whiskers show the full range across 200 alternative tie resolutions, not sampling CIs. **d,** Launch proportions after selecting 1,310 pairs by BRIDGE ranking or the local-transfer rule (265 and 223 launched pairs, respectively). Bars show proportions; whiskers show 95% indication-cluster bootstrap CIs. The bracket reports the paired difference in launch proportions (BRIDGE ranking minus local-transfer rule), in percentage points, with its 95% CI. Intervals use 5,000 resamples of 205 indication clusters, conditional on fixed selections. The top 572 empirical-*P* candidates constitute the *q* ≤ 0.05 support set; selecting 1,310 candidates is a ranking comparison. All CIs are two-sided. Source data are provided as a Source Data file.

A similar pattern was observed for the same-target transfer rule alone. This rule selected 510 pairs. Among these pairs, network-added pairs had a higher launch proportion than pairs not supported by BRIDGE (25.5% versus 14.5%; RS, 1.75; 95% CI, 1.20–2.55). BRIDGE also extended beyond same-target transfer, supporting an additional 282 pairs outside the rule, including 71 that had reached launch (**Supplementary Table 2**).

Because BRIDGE’s calibrated support set was smaller than the local-transfer selection, we examined whether the higher launch proportion persisted when the number of selected candidates was held equal. Specifically, we ranked all 7,763 non-seed candidates by increasing empirical *P* value and evaluated 11 selection cutoffs. Nine corresponded to the top 1–30% of candidates; two additional cutoffs selected 572 and 1,310 pairs, corresponding to the sizes of BRIDGE’s calibrated support set and the local-transfer rule’s selection, respectively (**Methods**). The observed launch proportions ranged from 16.3% to 29.7% across these cutoffs (**Fig. 3c**). At 10 of the 11 cutoffs, BRIDGE’s observed launch proportion exceeded the 17.0% obtained by the local-transfer rule. At a matched selection size of 1,310 pairs, BRIDGE included 265 launched pairs, compared with 223 under the local-transfer rule (20.2% versus 17.0%; difference, 3.21 percentage points; 95% CI, 1.12–4.97; **Fig. 3d**). Raw-score and z-score rankings also selected more launched pairs than the local-transfer rule at 1,310 pairs (**Supplementary Fig. 7d; Supplementary Table 2**).

### Temporal evaluation of BRIDGE support against subsequent clinical outcomes

Because BRIDGE’s genetic evidence, PPI network and disease-similarity data were derived from pre-2024 sources, we evaluated support against clinical outcomes recorded in 2024–2025. We matched T–I pairs by gene and indication between the 22 December 2022 baseline dataset and the 5 December 2025 Citeline snapshot. We retained pairs that had reached at least phase I and had no recorded launch before 2024. Unmatched pairs and launched pairs without a recorded launch year were excluded. The resulting analysis set, termed the risk set, comprised 6,183 pairs: 376 with direct genetic support, 345 with network-added support and 5,462 unprioritized pairs (**Fig. 4a**, **Supplementary Table 1** and **Methods**).

**Figure 4.**
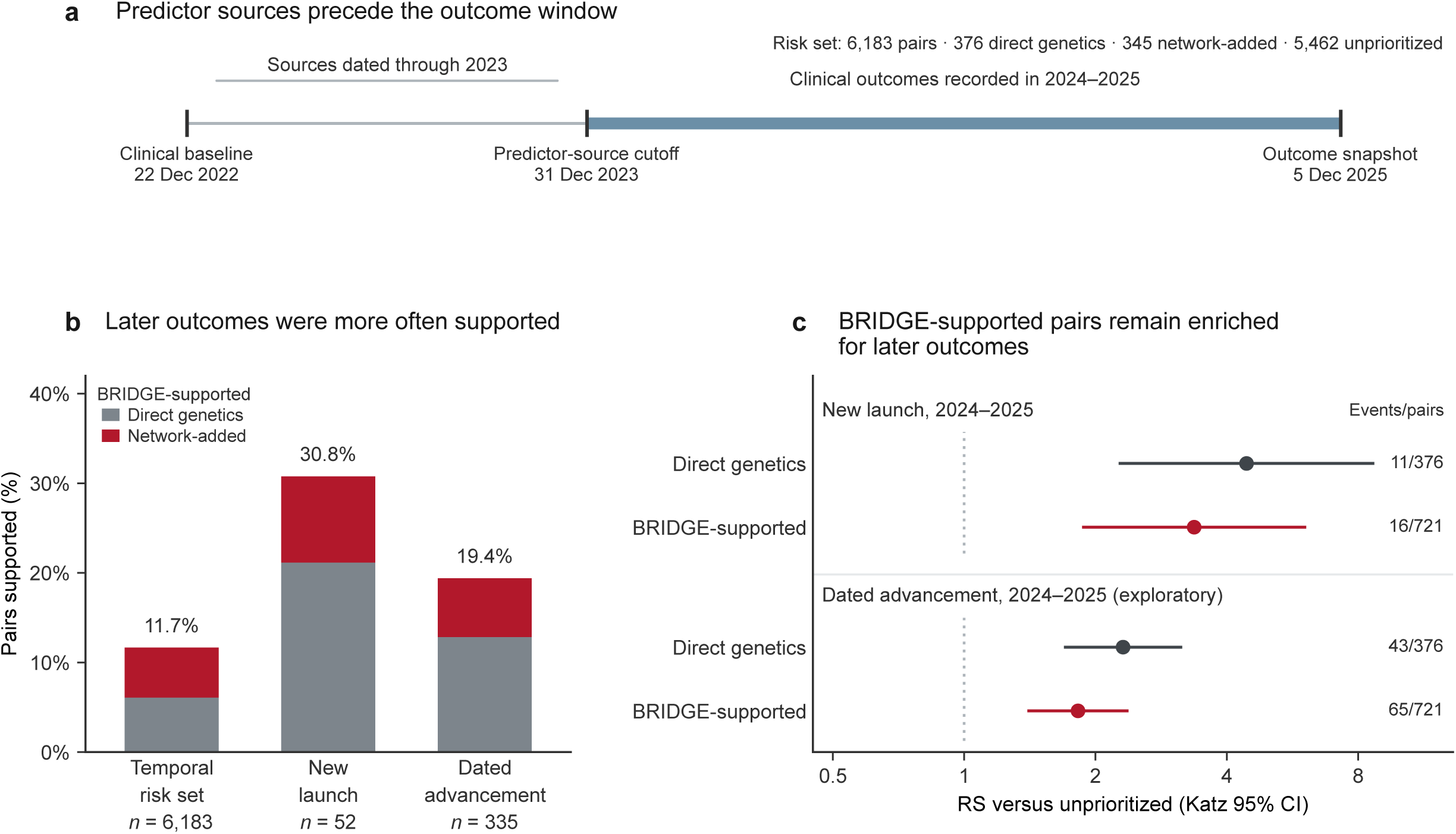
Temporal evaluation of BRIDGE support and clinical outcomes. **a,** Timeline of the 22 December 2022 clinical baseline, 31 December 2023 predictor-source cutoff and 5 December 2025 Citeline outcome snapshot. The risk set comprised 6,183 target–indication (T–I) pairs: 376 with direct genetic support, 345 network-added and 5,462 unprioritized. **b,** Proportions supported by full BRIDGE among all risk-set pairs (coverage), incident-launch pairs (launch capture) and dated-advancement pairs. Stacked bars distinguish direct genetic support (grey) and network-added support (crimson); labels give denominators and combined percentages. **c,** Relative success (RS) for direct genetic support and full BRIDGE support against the common unprioritized group. Incident launch required a recorded launch year of 2024 or 2025. Exploratory dated advancement combined a higher recorded stage than baseline with a most recent event date in 2024–2025; this date could concern another indication or an event without stage advancement (Methods; Supplementary Note 2). Filled circles and horizontal lines show RS and two-sided Katz 95% confidence intervals (CIs) on a logarithmic scale. Dark grey denotes direct genetic support and crimson denotes full BRIDGE; the dotted line marks RS = 1. Adjacent labels give events/pairs. Network-added estimates are given in **Supplementary Table 6**. Source data are provided as a Source Data file.

Full BRIDGE expanded support from 376 pairs with direct genetic support to 721 pairs. Among the 6,183-pair risk set, coverage increased from 6.1% to 11.7%. It supported 16 of 52 later launch pairs, compared with 11 with direct genetic support. Launch capture, the supported fraction of later launch pairs, therefore increased from 21.2% to 30.8% (**Fig. 4b**). The launch proportion used each support group as its denominator: 2.22% for full BRIDGE and 0.66% for unprioritized pairs (16/721 versus 36/5,462; RS, 3.37; 95% CI, 1.88–6.04; **Fig. 4c**). Network-added pairs had a launch proportion of 1.45% (5/345), compared with 0.66% for unprioritized pairs (36/5,462; **Supplementary Table 6**). The network-added estimate was exploratory and based on five launch pairs. Raw-record review showed that three of the five events came from a single anlotinib launch in nasopharyngeal cancer. A fourth, for KIT, came from the launch of a new imatinib formulation (**Supplementary Note 2** and **Supplementary Fig. 5b**).

We also examined dated clinical advancement as an exploratory endpoint. It combined a higher recorded stage than at baseline with a most recent recorded event in 2024–2025. This event date could concern another indication or an event other than the pair’s stage transition. Advancement rates were 9.0% for pairs supported by full BRIDGE (65/721), 6.4% for network-added pairs (22/345) and 4.9% for unprioritized pairs (270/5,462; **Supplementary Fig. 6a,b** and **Supplementary Table 6**).

### Genetic anchors connect the expanded target catalogue to therapeutic pathways

We next asked whether network propagation could nominate targets that could be altered by a therapeutic agent (therapeutically tractable targets) and that lack direct genetic support. To address this, we defined an expanded target catalogue of 101 genes, each with BRIDGE support but no direct genetic seed assignment in any indication. Together they account for 158 supported T–I pairs. To assess whether these targets could be pursued therapeutically, we matched them to the antibody and small-molecule tractability gene lists released by Minikel et al.^5^ These lists are derived from Open Targets and combine evidence from existing pharmacological agents with predictions of suitability for these treatment types. Of the 101 genes, 98 appeared on at least one list. For 26 of these 98 tractable genes, at least one supported pair had already reached launch in the Minikel et al. clinical dataset; the remaining 72 had none (**Fig. 5a**). Most catalogue genes were therefore tractable targets with no launched drug for their supported indications.

**Figure 5.**
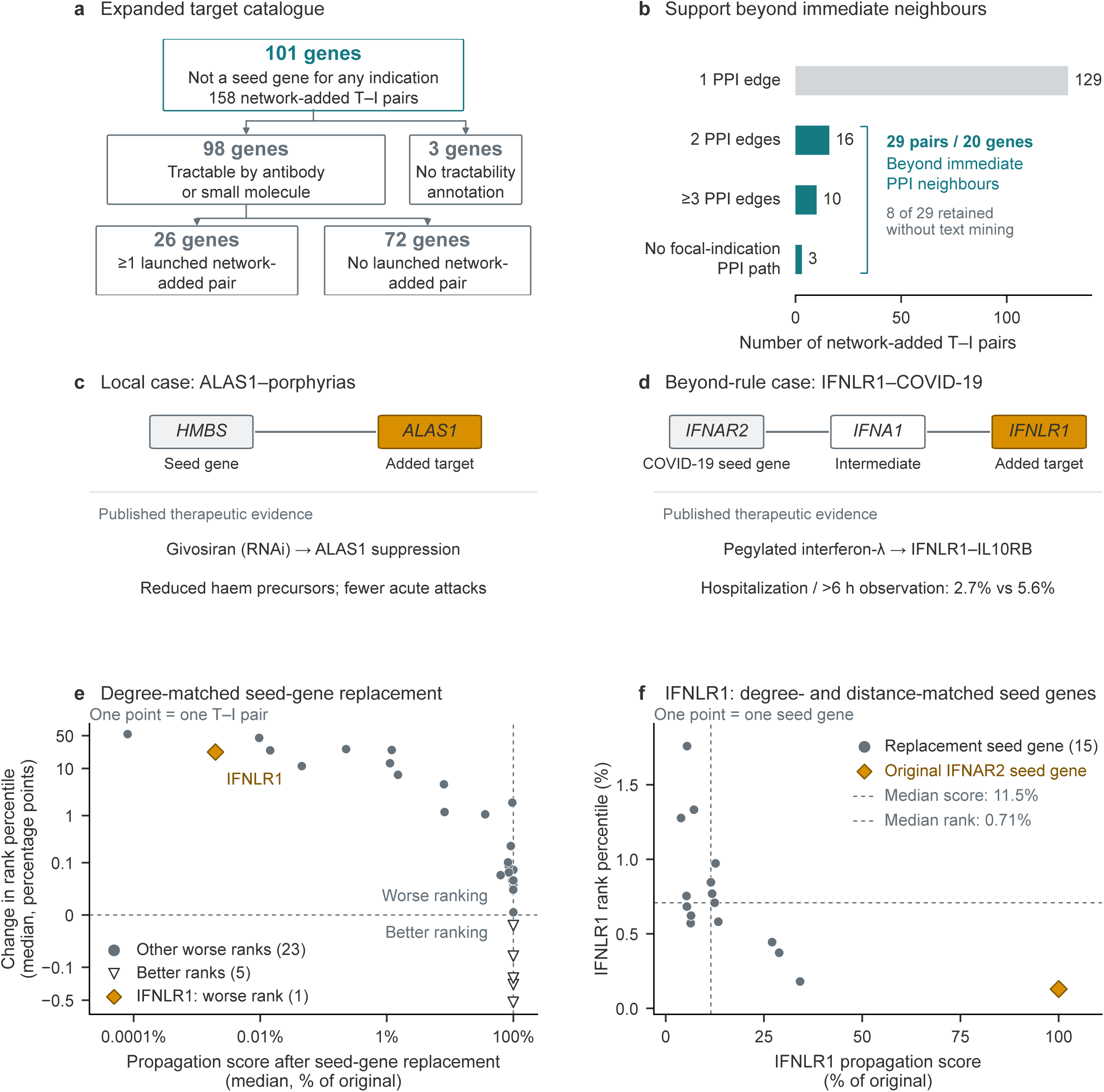
Therapeutic pathways and genetic-anchor sensitivity in the expanded target catalogue. **a,** The 101 genes with BRIDGE support but no direct genetic seed assignment in any indication, classified by antibody or small-molecule tractability annotations. For the 98 tractable genes, the lower split shows whether any supported target–indication (T–I) pair was launched in the baseline clinical dataset^5^. **b,** All 158 supported pairs classified by shortest unweighted protein–protein interaction (PPI) distance to a focal-indication seed. Teal bars identify the 29 pairs beyond immediate PPI neighbours. Support after text-mining exclusion was separately recalibrated. **c,d,** Retrospective examples of one-step ALAS1–porphyrias and two-step IFNLR1–COVID-19 support. Upper diagrams show selected undirected STRING functional associations, which can include indirect relationships. Grey, genetic seeds; white, intermediate; gold, added targets. *HMBS* evidence concerns acute intermittent porphyria, matched to porphyrias. Primary and no-text-mining *q* values are 0.03329 and 0.03134 for *ALAS1*, and 0.03349 and 0.04083 for *IFNLR1*. Lower diagrams summarize published evidence: givosiran reduced acute attacks in a randomized trial^24^; pegylated interferon-λ acts through IFNLR1–IL10RB^25^ and reduced 28-day hospitalization or emergency observation (>6 h) in a randomized trial (25/931 versus 57/1,018 placebo recipients)^26^. **e,** Anchor-replacement sensitivity for the 29 pairs beyond immediate PPI neighbours. Each point is one pair: median replacement score (% of original) versus median rank-percentile change across 100 degree-stratum-matched replacement configurations per indication. Axes are logarithmic (score) and symmetric-log (rank change; linear within ±0.1 percentage points). Grey circles, 23 other pairs with worse ranks; open triangles, five pairs with better ranks; gold diamond, *IFNLR1*–COVID-19. Rank change is replacement minus original (positive, worse). Dashed lines mark 100% retained score and zero rank change. **f,** *IFNLR1* scores and ranks with the original *IFNAR2* anchor and each of the 15 eligible replacements matched for weighted-degree stratum and two-step unweighted PPI distance. Each point is one seed assignment; the scored target is always *IFNLR1*. Grey circles, replacements; gold diamond, original anchor; dashed lines, replacement medians. Scores are percentages of the original; lower rank percentiles (among 19,699 genes) are better. Perturbation analyses summarize score and rank sensitivity with the original support assignments retained. Source data are provided as a Source Data file.

Next, we traced how each of the 158 pairs connected to genetic evidence for its focal indication. The target was one PPI edge from a focal-indication seed in 129 pairs, two edges away in 16 pairs and three or more edges away in 10. Three pairs had no PPI path to a focal-indication seed (**Fig. 5b**). The 29 pairs beyond immediate PPI neighbours, spanning 20 genes, therefore had neither direct genetic support for any indication nor a direct PPI link to a focal-indication seed. Their support arose through multi-step propagation across the gene and disease layers. To test whether this support depended on literature co-mention, we excluded STRING text-mining evidence. Eight of the 29 pairs remained supported, compared with 73 of the 129 directly connected pairs (**Supplementary Table 3**). Thus, BRIDGE extended genetic support beyond immediate neighbours even without text-mining evidence.

We selected two contrasting cases after scoring to examine the therapeutic interpretation of network-added candidates. Among directly connected pairs, ALAS1–porphyrias connects a disease-associated genetic anchor to an established therapeutic intervention at a pathway step preceding the inherited defect (**Fig. 5c**). The *HMBS* seed, derived from acute intermittent porphyria and matched to the broader porphyrias indication, was directly connected to *ALAS1* in the PPI network. Suppressing *ALAS1* in the liver reduces the accumulation of toxic haem precursors, as demonstrated in a disease model,^23^ and the RNA interference (RNAi) therapy givosiran, which lowers ALAS1 expression, reduced acute attacks in a randomized trial.^24^ This example links the inherited defect to an upstream therapeutic intervention and illustrates a therapeutic modality, or treatment type, beyond the catalogue’s antibody and small-molecule annotations.

Beyond immediate PPI neighbours, IFNLR1–COVID-19 provides a complementary case linking network-derived genetic support to a clinically investigated receptor pathway (**Fig. 5d**). *IFNLR1* was two PPI edges from *IFNAR2*, the sole COVID-19 seed in the benchmark, and retained support in both the primary analysis (*q* = 0.033) and the analysis excluding text-mining evidence (*q* = 0.041). Pegylated interferon-λ acts through the IFNLR1–IL10RB receptor complex, and published treatment-response data showed induction of interferon-stimulated genes in IFNLR1-expressing immune cells.^25^ In a randomized phase 3 trial, the composite endpoint of COVID-19-related hospitalization or emergency observation for more than 6 hours within 28 days occurred in 25 of 931 treated participants (2.7%) versus 57 of 1,018 placebo recipients (5.6%; relative risk, 0.49; 95% Bayesian credible interval, 0.30–0.76).^26^ An earlier randomized trial found no significant reduction in viral shedding or symptom duration (**Supplementary Table 3**). These trials provide endpoint-specific therapeutic evidence for the IFNLR1–IL10RB receptor pathway linked to the network-added candidate.

We next asked whether BRIDGE’s candidate scores and ranks depended on which genes were used as genetic anchors. For all 29 pairs beyond immediate PPI neighbours, we replaced the focal-indication seed genes with genes from the same weighted-degree strata, transferring each seed gene’s assignments to its replacement (Methods). Median propagation scores were lower than with the original seeds for every pair; median ranks worsened for 24 and improved for five (**Fig. 5e**; **Methods** and **Supplementary Table 3**). A lower score can still yield a better rank if other genes’ scores fall more.

For *IFNLR1*–COVID-19, we then replaced *IFNAR2*, the sole COVID-19 seed, with each of 15 eligible genes. All replacements matched *IFNAR2*’s weighted-degree stratum and its shortest unweighted PPI distance of two steps from *IFNLR1*. This matched broad connectivity and distance to the target. Across these replacements, the median propagation score was 11.5% of the score obtained with *IFNAR2*. Gene-rank percentiles place IFNLR1 among all 19,699 genes, with lower values indicating better ranks. The median gene-rank percentile was 0.71% (range across replacements, 0.18–1.76%), compared with 0.13% with *IFNAR2* (**Fig. 5f**). Using *IFNAR2* as the anchor therefore gave *IFNLR1* a better rank than every matched replacement, although all replacements retained it within the top 1.8% of genes.

Together, the catalogue and case studies connect network-derived genetic support to clinically investigated therapeutic pathways, including candidates beyond immediate PPI neighbours of genetic seeds. Connecting these candidates to identifiable genetic anchors, assessing their sensitivity to anchor replacement and integrating published pharmacological evidence provides a traceable basis for further target- and indication-specific evaluation.

## Discussion

BRIDGE expands clinically informative genetic support for therapeutic target prioritization by coupling disease relationships with protein connectivity. In our retrospective evaluation of the Minikel *et al.* benchmark, combining direct genetic support and network-added support nearly doubled coverage while retaining enrichment for launches.^5^ The expansion reached additional indications for seed genes and 101 genes outside the benchmark’s direct genetic seed set. These findings extend the established association between direct genetic support and development outcomes to additional T–I hypotheses.^3,5^

These findings build on approaches that use molecular networks to interpret genetic associations^13–16^ and related-trait evidence to recover approved drug targets.^27^ Disease relationships provide access to direct genetic support for the same target in related indications, while protein connectivity extends that evidence to other targets. Withholding and restoration experiments separated these routes and linked recovery to the availability of relevant genetic anchors. Coupling the two therefore allows genetic evidence to inform additional intervention points across related disease settings.

The local-transfer comparisons show that network integration provides additional information for prioritization. Most network-added pairs already had local connections through related indications or PPI neighbours. BRIDGE distinguished clinically enriched subsets within these candidates and identified another enriched subset outside the rule. Its empirical-*P* ranking captured more launched pairs when the number selected matched the local-transfer rule’s selection size. This additional discrimination provides a basis for selecting candidates for further evaluation, including those that simple local rules omit.

Pair-specific empirical calibration translates propagation scores into an explicit support criterion. Building on work on network connectivity and score normalization,^19,20^ BRIDGE compares each candidate’s observed score with scores for the same pair after degree-matched seed replacement. A common threshold after adjustment for multiple testing defines support without fixing the catalogue size in advance. For 572 selected pairs, empirical-*P* and raw-score rankings captured equal numbers of launched pairs, while empirical-*P* ranking selected less connected targets (**Supplementary Fig. 7a–c**). Random-seed diagnostics without designated true signals produced no support calls.

The expanded catalogue provides a traceable connection between genetic evidence and therapeutically relevant intervention points. The retrospectively selected *ALAS1* and *IFNLR1* cases illustrate complementary uses of this information. *ALAS1* connects an inherited metabolic defect to an upstream intervention, whereas *IFNLR1* links a target beyond immediate PPI neighbours to a clinically investigated receptor pathway.^23–26^ For *IFNLR1*, the original *IFNAR2* seed improved ranking relative to matched alternatives, which also ranked the target highly. This pattern shows how a specific genetic anchor can refine prioritization within a broader network context. Pharmacological evidence adds intervention direction, modality and disease context, including the IFNLR1–IL10RB receptor complex for *IFNLR1*. Tractability annotations and development records provide further context for selecting indication-specific follow-up (**Supplementary Fig. 9**).

The retrospective temporal analysis extended the coverage finding to launches recorded after the dates of the input sources. Full BRIDGE captured more later launch pairs than direct genetic support alone.

There are several limitations to this study. Seed membership depends on locus-to-gene mapping and trait–indication matching, so genes outside the seed set may still have relevant genetic associations. Functional associations and disease similarity lack tissue-specific context, and research attention can influence the underlying networks.^17,21^ Calibrated support was assessed in seeded indications, whereas cross-indication recovery evaluated gene rankings using neighbouring seeds. When artificial score increases were assigned to 0.1–1% of candidates, none with increased scores received support. General false discovery rate control and biological sensitivity require further validation (**Supplementary Note 1**). The benchmark comprised pairs already in human development and included dependent outcomes from shared genes, indications and programmes. The network-added estimate rested on five launch pairs, three from one programme, limiting precision. Joint stratification by baseline activity, phase and indication yielded an imprecise full-BRIDGE temporal estimate (**Supplementary Note 2**). Longer follow-up across independent programmes with documented evidence dates would clarify the magnitude and consistency of network-added enrichment.^29^ Disease-relevant experiments are needed to establish causal mechanisms and intervention direction.^6,28^

BRIDGE complements direct genetic support with traceable network-derived hypotheses across targets and indications. Combining these hypotheses with mechanistic and pharmacological evidence provides a practical basis for selecting candidates for prospective evaluation. Studies with predictions and comparators fixed before follow-up can test whether this broader genetic support improves target–indication decisions.

## METHODS

Methods are available in the **Methods** section.

## ACKNOWLEDGEMENTS

Research reported in this work was supported by National Cancer Institute under Award Numbers R01CA263494, U01CA293883, and P30CA016672; and The U Foundation U-Pilot Award. The content is solely the responsibility of the authors and does not necessarily represent the official views of the National Institutes of Health.

## Methods

### Study design and genetic evidence

We analysed T–I pairs from the Citeline Pharmaprojects cohort and genetic-evidence tables released by Minikel *et al.*^5^. Eligibility required a known outcome for at least one transition from phase I to II, phase II to III or phase III to launch. Indications also required the source’s genetic-insight annotation, based on trait–indication similarity ≥0.8. Qualifying evidence came from Online Mendelian Inheritance in Man (OMIM), IntOGen or a genome-wide association study (GWAS) trait with at least three independent loci^5^. The released genetic-insight annotation determined indication eligibility. Among 13,030 eligible rows, eight T–I identifiers were duplicated. We retained the highest combined maximum development phase, then ranked transition outcomes from phase III→launch, phase II→III, phase I→II and pre-phase I→phase I. Within each transition, success took precedence over failure, then unknown outcome; remaining ties followed source order. This yielded 13,022 unique pairs across 670 indications (**Supplementary Table 1**). The analysis included all eligible pairs in the released cohort.

Direct genetic support required normalized trait–indication similarity ≥0.8 and qualifying OMIM^30^, Open Targets Genetics, PICCOLO expression quantitative trait locus (eQTL) or Genebass evidence^31^. Open Targets Genetics required locus-to-gene share ≥0.5^5,8^; PICCOLO required colocalization posterior probability H4 ≥0.9^5^. Genebass included burden and sequence kernel association tests. The released association calls defined the qualifying genetic evidence. Somatic evidence could qualify an indication for inclusion but was excluded from seeds. Of 667 pairs with direct genetic support, 658 mapped by exact gene symbol to STRING v12.0^21^ and became seeds. The nine unmapped pairs remained in whole-cohort analyses. Network evaluation comprised 658 seeds and 7,763 non-seed candidates across 241 seeded indications. Gene-level seed membership was defined across the 658 assignments, comprising 359 genes, before evaluating network support or outcomes.

The study used a retrospective clinical benchmark that had also informed earlier method iterations. Following cohort deduplication, network construction, scoring and support assignment used the genetic-evidence and network inputs. Clinical outcomes were joined after support assignments were fixed.

### Network construction and propagation

The PPI layer retained STRING v12.0 associations with combined confidence ≥0.90, weighted by score/1,000^21^. Reciprocal records were combined by maximum weight, yielding 100,856 undirected edges among 19,699 mapped genes, of which 12,323 were non-isolated. Unweighted degree counted neighbours; weighted degree summed incident weights. The released MeSH-similarity matrix^5^ covered 669 indications represented by mapped cohort pairs. Retaining similarities ≥0.50, removing self-edges and taking maximum reciprocal weights yielded 9,205 disease edges. The 658 seed pairs supplied unit-weight gene–disease links.

Propagation used a coupled random walk with restart^13^, with a state containing both gene and disease scores. Within-layer and cross-layer operators were normalized separately over outgoing weights. Zero-sum columns remained zero; unavailable transition weights were not redistributed. Starting from the focal-indication unit vector *p*^(0)^ = *e_d_*, the update was

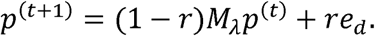

The operator *M_λ_* weighted within-layer and cross-layer transitions by 1 – *λ* and *λ*, respectively, without global renormalization. Primary restart and coupling probabilities were *r* = 0.50 and *λ* = 0.50. Iteration stopped when the combined gene- and disease-layer *ℓ*_1_ change was below 10^-7^ for every indication, or after 300 iterations. The final gene-layer value was the propagation score for pair (*g,d*). Numerical settings specific to target withholding and anchor restoration are given below.

### Calibration and support assignment

The 12,323 non-isolated genes were divided into ten approximately equal-sized pools by weighted PPI degree. Each seed assignment defined a disease-labelled slot. Replacement genes were drawn independently and uniformly, with replacement, from the original seed’s pool. Four isolated seed slots used the lowest non-zero-degree pool. Repeated replacements retained multiplicity in gene–disease weights. Graphs, walk parameters and each indication’s seed-slot counts and degree-pool composition remained fixed; gene–disease transitions and propagation were recomputed.

Three independent blocks each contained 2,100 resamples: 2,000 reference and 100 held-out configurations. The pooled 6,000 references defined each candidate’s null distribution, including configurations in which it became a focal-indication seed. The 300 held-out configurations were reserved for diagnostics. Reference scores were stored in float32, and observed scores were cast to the same precision before comparison. For candidate *i*,

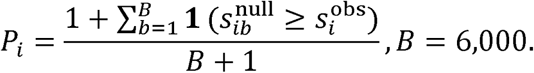

Here, 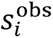 and 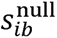 denote observed and resampled scores. Candidates shared resampling configurations but were compared only with their own null scores. Ties counted as exceedances, and the plus-one correction gave a minimum probability of 1/6,001^32^. BH adjustment^22^ was applied jointly across the 7,763 non-seed candidates; *q* ≤ 0.05 defined network-added support. Direct seeds were excluded from testing. Full BRIDGE combined direct genetic support and network-added support. Degree-pool cutpoints and seed-slot allocations are provided in **Source Data, degree-pool specifications. A**lternative reference definitions and blockwise diagnostics are described in **Supplementary Note 1**.

### Clinical evaluation and sensitivity analyses

Coverage was the supported fraction of the specified cohort or clinical-status group. Resolved historical status took precedence over active status for mutually exclusive groups. RS was the event proportion in a focal set divided by that in its comparator. For whole-cohort launch analyses, direct genetic support and full BRIDGE each used their own complement among 13,022 pairs. Network-added support used the 7,191 unprioritized eligible candidates. Within-stratum comparisons used the unprioritized remainder of that stratum. Katz log-scale intervals for RS used no continuity correction^33^; Wilson intervals described event proportions and coverage^34^. Clinical comparisons used unadjusted tests and nominal 95% confidence intervals; intervals were two-sided unless specified otherwise. Analytic intervals treated pairs as independent; cluster-based comparisons are specified below.

Therapy-area analyses used the released many-to-many map to 17 MeSH areas^5^, counting each pair once per area. Development-transition analyses excluded unknown outcomes separately for each transition. Six genetic-evidence sources were calibrated separately: OMIM, Genebass, pooled genetics, Open Targets Genetics GWAS, augmented Open Targets genetics and PICCOLO eQTL. The pooled source combined the four qualifying sources used for primary direct genetic support. The augmented Open Targets source used all 864 seed pairs in its fixed ledger. Each source had its own seeds, candidate family, 6,000 reference configurations and 300 held-out configurations, with candidate testing and multiplicity adjustment performed within each source. All sources used STRING confidence ≥0.90. Raw-score ranking frontiers were compared with degree-matched resample rankings (**Source Data, six-source ranking frontiers**).

One-factor-at-a-time sensitivities varied STRING confidence (0.40, 0.70, 0.90) and disease similarity (0.30, 0.50, 0.70, 0.90). They also varied restart probability (0.30, 0.50, 0.70) and inter-layer coupling (0.25, 0.50, 0.75). Each network or walk variant received its own 6,000 reference and 300 held-out configurations. Support-threshold sensitivities at *q* = 0.01, 0.02, 0.05, 0.10 and 0.20 re-used the primary adjusted probabilities (**Supplementary Fig. 2**; **Supplementary Tables 1 and 2**).

A separate sensitivity reconstructed STRING confidence without text-mining evidence^21^, using neighbourhood, fusion, co-occurrence, co-expression, experimental and curated-database channels. For each channel score *S_c_* scaled to 0–1, we removed the prior *π* = 0.041 as 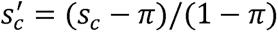. Negative adjusted scores were set to zero before combining channels. We combined channels and restored the prior: 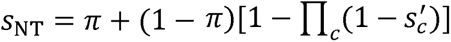 Edges with reconstructed confidence ≥0.90 formed the sensitivity graph, which was recalibrated using 6,000 new references and 300 held-out configurations.

### Disease-context and target-recovery analyses

Architecture comparisons used PPI-only propagation, a coupled walk without disease–disease edges and the full network. Each used the same seeds and candidates with separate calibration. PPI-only propagation restarted uniformly over the focal indication’s seeds. One hundred global disease-label shuffles applied the same permutation to rows and columns of the 669-node disease adjacency. This preserved weighted topology while disrupting alignment with genetic evidence; each shuffled network was recalibrated. Upper-tail empirical probabilities were (1 + *k*)/101, where *k* counted shuffled statistics at least as large as observed. Matched-depth raw-score comparisons selected 572 candidates per architecture or shuffle, independently of calibrated support. Additional ranking-depth comparisons accompany **Supplementary Table 2**; shuffle-overlap diagnostics are described in the caption to **Supplementary Fig. 3**.

Unweighted shortest paths measured molecular distance from a target to a focal-indication seed. Disease-context distance connected the focal indication to an indication seeding the same gene, using disease edges with MeSH similarity ≥0.50. Distances were computed before joining support or outcomes, and unreachable pairs remained in the analysis. Distance categories are defined in **Fig. 2d**.

Target withholding assessed rank recovery independently of calibrated support calling, with a rank percentile ≤5% defining recovery. PPI-only leave-one-target-out analysis included 139 launched pairs with direct genetic support across 66 indications. Each withheld target and at least one remaining focal-indication seed had to be non-isolated. The target remained an unlabelled node; propagation restarted uniformly over remaining non-isolated seeds with r = 0.50. These walks used an *ℓ*_1_ tolerance of 10^-8^ and at most 300 iterations. Propagation and unweighted degree ranked the same non-isolated genes, excluding remaining seeds. For average descending rank *R* among *N* eligible genes, the rank percentile was 100(*R* – 1)*N*. A one-sided Wilcoxon signed-rank test^35^ assessed whether propagation gave lower target-rank percentiles (**Supplementary Fig. 4a** and **Supplementary Table 4**).

Leave-one-disease-out analysis removed all focal-indication seed assignments. Eligibility required a mapped launched pair with direct genetic support and a seeded neighbouring indication, yielding 181 pairs across 108 indications. The primary cross-target cohort comprised 49 pairs whose targets had no remaining seed assignment; the other 132 retained same-target evidence elsewhere. The full walk was compared with a control replacing the PPI transition matrix by the identity, retaining disease and gene–disease transitions. Targets were ranked among all 19,699 mapped genes. Rank percentiles were 100(*n*_>_ + *n*_=_/2)/*N*, where *n*_>_ and *n*_=_ counted genes scoring above and equal to the target, including the target among ties. Walks used the primary tolerance of 10^-7^ and at most 300 iterations. One thousand global disease-label permutations shared a row-and-column permutation across focal indications, with fixed eligibility. Empirical probabilities were (1+*k*)/1,001; recovery proportions used Wilson intervals (**Supplementary Table 4**). These permutations were separate from the 100 recalibrated shuffles.

Anchor-restoration cohorts and perturbations were specified after inspection of withholding results but before new scoring. Real anchors were distinct genes seeded in immediately neighbouring indications. The count experiment retained 39 cross-target pairs across 33 indications with at least eight anchors. Nested random subsets restored 0, 1, 2, 4 or 8 anchors across 50 shared backgrounds per indication. Inactive anchors received unique surrogates from the same weighted-degree stratum, with isolated genes kept separate. Surrogates excluded all original seed genes and the 49 held-out targets. Every occurrence of an inactive anchor received the same surrogate; restoration returned all of that gene’s seed assignments. This preserved seed-assignment counts, degree-stratum composition, distinct seed-gene counts, multiplicities and cross-indication sharing. Focal-indication seeds remained withheld and targets remained unseeded elsewhere; zero restored anchors retained the surrogate background.

Fractional restoration included all 49 eligible pairs across 41 indications, restoring 0%, 25%, 50%, 75% or 100% of available anchors. Nonzero anchor counts were rounded upwards, so equal fractions could restore different numbers of genes (**Supplementary Fig. 4b**). A further sensitivity retained the nearest real anchor while restoring 1, 2, 4 or 8 anchors in the 39-pair count-restoration cohort. This preserved the minimum distance to a real anchor across restoration levels (**Supplementary Fig. 4c**). Additional recovery thresholds assessed dependence on the ranking cutoff (**Supplementary Fig. 4d**; **Supplementary Table 4**).

Gene–disease transitions were recomputed after each perturbation. Restoration used *r* = *λ* = 0.50, an *ℓ*_1_ tolerance of 10^-11^ and at most 1,500 iterations. Targets were ranked among 19,699 genes using the disease-withholding midpoint convention. Recovery was averaged over backgrounds within pairs, then over pairs. Paired changes and pointwise intervals used 5,000 focal-indication cluster resamples, conditional on the sampled backgrounds. The backgrounds represented Monte Carlo seed configurations, and the analysis summarized rank recovery of known positive pairs.

### Local transfer and normalization comparisons

Same-target transfer required direct genetic support for the candidate gene in another indication with MeSH similarity ≥0.50. The local-transfer rule selected the union of this group and targets one PPI edge from a focal-indication seed, yielding 1,310 candidates. Within-rule comparisons used the unprioritized remainder of the same stratum.

The 7,763 candidates were ranked by ascending empirical *P*, descending raw propagation score or descending null z score^20^. For *z_i_* = (*s_i_* – *μ_i_*)/*σ_i_*, *μ_i_* and *σ_i_* were the mean and sample standard deviation (SD) of candidate’s 6,000 reference scores. Diagnostic configurations were excluded from this calculation. When *σ_i_* = 0, positive, zero and negative deviations ranked at +∞, 0 and -∞, respectively. Z scores were used only for ranking, and primary ties followed ascending T–I identifier. A sensitivity assigned all candidates with zero reference variance the minimum finite z score, then repeated the ranking comparisons. Nine percentage depths (1%, 2%, 3%, 5%, 7.5%, 10%, 15%, 20% and 30%) used half-up rounding of candidate counts. Two additional depths selected exactly 572 and 1,310 pairs. The top 572 empirical-*P* candidates equalled the calibrated support set; other depths were ranking comparisons.

Comparisons with the local-transfer rule in **Fig. 3** used 5,000 shared indication-cluster bootstrap resamples across the 205 indications represented among candidates. These resamples estimated launch proportions, within-stratum RS and paired differences in launch proportions. Matched-ranking comparisons and launch capture used the same bootstrap approach, with selections fixed and denominators recalculated within each resample. Launch capture used all 765 launched candidates as its denominator. Intervals were pointwise percentile intervals, conditional on fixed scores, references and selections. The same-target-only comparison in **Supplementary Table 2** instead used Wilson intervals for launch proportions and Katz intervals for RS. Matched-depth RS used each ranking’s own unselected remainder and Katz intervals.

A separate bootstrap resampled genes as clusters to assess sensitivity to shared genes. Two hundred random identifier orders shared across methods and depths assessed tie sensitivity. Scores and identifier order determined each ranking. These comparisons were exploratory. Additional normalization and ranking results accompany **Supplementary Table 2** and **Supplementary Fig. 7**.

### Assessment of calibration

Complete-null diagnostics used held-out configurations as pseudo-observed candidate families and repeated empirical testing and BH adjustment. N0 used the primary independent degree-matched replacement; N1 replaced isolated seeds from a separate isolated-gene pool. N2 applied a shared one-to-one gene replacement within N1 pools, preserving seed-gene multiplicities and cross-indication sharing. Each model supplied 6,000 references and 300 held-out configurations. The primary diagnostic retained the original candidate family, excluding candidates seeded for their focal indication in the held-out configuration. Empirical probabilities used all 6,000 unfiltered references. Every selection under a complete null counted as false. Any-call frequencies received one-sided 95% Clopper–Pearson upper bounds^36^ (**Supplementary Fig. 8a** and **Supplementary Table 5**).

The primary score-shift experiment used N0 scores to assess recovery and false selections under artificial increases (**Supplementary Fig. 8b,c**). Candidate-specific sample standard deviations (SDs) were estimated from the first block’s 100 held-out pilot configurations. The other 200 configurations were evaluated against all 6,000 references, retaining the original candidates except focal-indication pseudo-seeds. Nested candidate sets were sampled within weighted-degree strata in approximate proportion to eligible stratum sizes, at prevalences of 0.1%, 0.5%, 1%, 5% and 10%, with total counts rounded half-up. Scores were increased by 0.5, 1, 2, 3, 4, 6, 8 or 12 pilot SDs. The same candidate sets were used across strengths and comparator procedures. Shifts were applied in float64 without rounding back to float32.

Only candidates whose scores strictly increased counted as effective simulated signals. Recall was the fraction of these candidates recovered and was undefined when no effective signals existed. The false discovery proportion (FDP) was the fraction of calls without an effective increase, set to zero when there were no calls; designated candidates whose scores did not increase remained null for FDP. Mean recall averaged families with at least one effective signal; mean FDP and no-call frequency used all 200 evaluation families. Each contributing family received equal weight. Pointwise percentile intervals used 5,000 paired family-bootstrap resamples, conditional on references and pilot scales. Bootstrap means likewise excluded undefined recall values. Score perturbations were applied to stored propagation outputs, and performance was evaluated under the specified score-shift models. Reference-size and eligibility sensitivities, N2 experiments, alternative scales and comparator procedures are in **Supplementary Note 1** and Source Data. Historical 4,000-reference diagnostics are summarized in **Supplementary Table 5**; blockwise support stability is shown in **Supplementary Fig. 8d**.

### Temporal evaluation of clinical outcomes

Retrospective temporal evaluation compared the 22 December 2022 clinical dataset with the 5 December 2025 Citeline snapshot. Genetic and network inputs comprised the released genetic and disease-similarity tables^5^ and STRING v12.0^21^. Scores and support labels were fixed before follow-up outcomes were joined. Matching normalized gene symbols and MeSH identifiers linked 7,049 of 8,421 primary-context pairs. We excluded unmatched pairs, baseline launches, launches recorded before 2024 and launched pairs without a recorded launch year. The resulting risk set contained 6,183 pairs (**Supplementary Table 1**).

The primary endpoint required a launched follow-up status and a recorded launch year of 2024 or 2025. Exploratory dated advancement required a higher stage than baseline and a most recent recorded event date in 2024–2025. Ordered stages were phase I, phase II, phase III, pre-registration/registration and launch. Advancement was introduced after preliminary arm-level feasibility review and before the amended analysis. The date field described the most recent recorded development event, which could relate to another indication or to an event other than a stage transition. Dated advancement therefore combined observed stage change with this event-year annotation. The five network-added launch pairs were checked against indication-specific raw records, as detailed in **Supplementary Note 2**.

Primary support-arm comparisons used the common unprioritized group, Wilson event-rate intervals, Katz RS intervals and two-sided Fisher exact tests. Mantel–Haenszel odds ratios^37^ stratified by baseline phase, indication or both, retaining strata containing both comparison groups without zero-cell correction (**Supplementary Fig. 5a**). Exploratory activity analyses used a nonmissing active-status field (acat) as the recorded-activity indicator; missing values represented unrecorded activity status. Stratification used activity alone, activity with phase, and activity with phase and indication. These analyses and the raw-score comparison were added after inspection of initial temporal results. Raw-score selection was fixed before temporal filtering, and each selection rule was compared with its own non-seed remainder. Additional sensitivities used 5,000 indication-cluster bootstrap resamples and omission of event-containing indications (**Supplementary Fig. 5a,b**). Indication-stratified conditional logistic models are reported in **Supplementary Table 6**. Model specifications, alternative advancement definitions and raw-score selection details are provided in **Supplementary Note 2**; estimates are in **Supplementary Table 6**.

### Target catalogue and genetic anchors

Added genes had network-added pairs but no assignment in the 359-gene mapped seed set. Antibody and small-molecule annotations were assigned by matching gene symbols to the gene lists released by Minikel et al.^5^ Derived from Open Targets, these lists combined pharmacological precedence with predicted tractability. A gene counted as annotated if it appeared in either list. Within the supported catalogue, a gene counted as launched if any of its BRIDGE-supported pairs was recorded as launched in the Minikel et al. clinical dataset. Development elsewhere was the maximum recorded phase across other indications in the 13,022-pair cohort, excluding the focal indication. Categories were assigned after support calling and could differ across focal indications for the same gene. Comparisons used the unprioritized remainder within each development category (**Supplementary Fig. 9**).

For all 5,463 candidates outside the seed gene set, we summarized uncapped shortest PPI distances to focal-indication seeds and connections to neighbouring seeded indications. Unreachable candidates formed a separate category. For the 158 supported pairs, we recorded directly connected seeds, all tied nearest focal-indication seeds and one deterministic representative shortest route through the full network. Distances used unweighted retained edges; full-network routes could traverse PPI, gene–disease and disease–disease edges. These routes recorded the network connections linking candidates to genetic seeds (**Supplementary Table 3**).

Within the 158 supported pairs outside the seed gene set, the combined local-transfer rule reduced to a direct PPI connection to a focal-indication seed. We cross-checked the resulting 29 beyond-rule pairs against the full local-rule ledger and summarized retention of support after text-mining exclusion (**Supplementary Table 3**).

An exploratory seed-replacement analysis included all 29 beyond-rule pairs across 26 indications. Its protocol was fixed after inspection of existing calls, paths and case literature but before new scoring. For each indication, all focal-indication seed genes were jointly replaced in 100 configurations by unique non-seed genes from the same weighted-PPI-degree strata. The replacement pool excluded the 20 candidate genes. Each replacement gene took over every assignment of its corresponding seed, including assignments in other indications. This preserved seed-assignment counts, degree-stratum composition, gene multiplicities and cross-indication sharing. Other seed assignments, PPI and disease–disease edges, and walk parameters remained fixed; gene–disease transitions were renormalized. Before perturbation, we verified reproduction of all 8,421 original observed scores. Original and replacement scores were then computed using an ℓ₁ convergence tolerance of 10^−11^ and at most 1,500 iterations. Rank percentiles among all 19,699 genes used the disease-withholding convention. We summarized computational sensitivity using pair-specific medians and 5th–95th percentile ranges across replacement configurations (**Supplementary Table 3** and accompanying Source Data).

For *IFNLR1*–COVID-19, *IFNAR2* had one seed assignment globally, so global and focal-indication replacement were identical. Because matching on weighted-PPI-degree stratum alone did not preserve the distance to *IFNLR1*, we performed a second exploratory comparison additionally matching shortest unweighted PPI distance. The protocol was fixed before scoring all 15 eligible replacement genes in the same weighted-degree stratum and at the same two-step distance. We summarized *IFNLR1* propagation scores and rank percentiles using the median and full range across this complete set of matched replacements (**Supplementary Table 3**).

Therapeutic examples were selected retrospectively after scoring for traceable genetic seeds and published target-specific or receptor-pathway evidence. *ALAS1*–porphyrias illustrated a connection within the local-transfer rule. *IFNLR1*–COVID-19 illustrated a connection outside the rule that retained support after text-mining exclusion. We checked original source traits, indication matches, PPI edges and results from the separately recalibrated analysis excluding text-mining evidence. Experimental and clinical reports supplied the intervention direction, modality and disease-subtype context for interpreting the fixed computational results. The full beyond-rule catalogue, original seed-source records, prior phase III status and positive and negative intervention evidence accompany **Supplementary Table 3**.

### Software and reproducibility

Analyses used Python, NumPy^38^, SciPy^39^, pandas^40^, statsmodels^41^ and Matplotlib^42^. Random seeds and computational provenance are recorded with the analysis outputs. Analyses used T–I records without participant-level data.

### Data availability

Source data are provided in the accompanying Source Data file. The processed data will be made publicly available on GitHub and Figshare upon publication.

### Code availability

Codes will be publicly available on GitHub upon publication. For peer review, the code is provided in BRIDGE_Reproducibility.zip, together with processed model inputs, software requirements and instructions for reproducing the analyses and figures.

### Author contributions

W.Z. and C.W. designed the study. W.Z. and C.W. developed the methods. W.Z. analyzed the data and wrote the initial version of the manuscript. J.X, Y.L, R.S., H.S., L.W., B.Z., and HW.D. provided critical feedback, improved the overall study design, and contributed to the interpretation of the results. All authors revised the manuscript.

