## Supplementary Figure 1; Supplementary Note 1 for "Integrating disease and protein networks expands genetic support for therapeutic targets"

Supplementary Information

**Supplementary Tables 1–6 are supplied in the accompanying Excel workbook. Numerical records and code for figure and table analyses accompany the BRIDGE_Reproducibility.zip archive.**

### Supplementary Notes

These notes extend the primary procedures in **Methods** with calibration and temporal sensitivity analyses. BRIDGE denotes Biological Relation–Informed Diffusion of Genetic Evidence. Supplementary Figures 1–9 and Tables 1–6 provide detailed results; case sources accompany **Supplementary Table 3**. Analyses used target–indication (T–I) pairs, each combining a target gene with a disease or condition considered for treatment, without participant-level data. The focal indication is the indication currently being evaluated. Direct support denotes qualifying genetic evidence matched to that indication; mapped directly supported pairs provide the network seeds. Network-added pairs lack direct support for the focal indication but meet BRIDGE’s calibrated support threshold. BRIDGE-supported pairs combine direct and network-added support. Confidence intervals (CIs) are two-sided 95% unless otherwise stated.

#### Supplementary Note 1. Sensitivity of empirical calibration

We examined sensitivity to the null models defined in **Methods**, Assessment of calibration: independent degree-matched replacement (N0), a separate isolated-gene pool (N1), and one-to-one replacement preserving seed-gene reuse (N2). N2 retained 359 distinct seed genes, their multiplicities and cross-indication sharing, up to relabelling. The three models produced similar support counts but different membership (**Supplementary Table 5**). The only N0-supported pair with zero reference variance, *DPEP1*–D000086382, remained supported under N1 and N2 with positive variance. Separately, we removed reference configurations in which the candidate became a focal-indication seed gene. Empirical probabilities then used candidate-specific reference counts $B_{i}$ and denominator $B_{i}+1$^1^. Reference filtering changes the null comparison; it is distinct from excluding candidates from the pseudo-observed testing family.

Complete-null diagnostics crossed the candidate and reference-filtering choices below with two reference sizes, yielding 14 conditions per null model. The 6,000-reference analysis pooled all three blocks; the 4,000-reference analysis omitted the held-out configuration’s block. Empirical testing and Benjamini–Hochberg (BH) adjustment^2^ were repeated within each eligible candidate family.

| **Starting candidate family** | **Candidates excluded from the pseudo-observed family** | **Reference configurations** |
| --- | --- | --- |
| All 8,421 mapped pairs | None | All configurations |
| All 8,421 mapped pairs | Candidates drawn as focal-indication seed genes | All configurations, or configurations with focal-indication seeding removed |
| Original 7,763 candidates | Candidates drawn as focal-indication seed genes | All configurations, or configurations with focal-indication seeding removed |
| Original 7,763 candidates | Genes seeded in any indication | All configurations, or configurations with focal-indication seeding removed |

No condition produced calls in any of its 300 held-out families. The one-sided Clopper–Pearson 95% upper bound on the frequency of any call was 0.994%^3^ (**Supplementary Fig. 8a**; **Supplementary Table 5**). Conditions within each model reused held-out families and were not independent replications. An earlier 4,000-reference diagnostic excluded genes seeded anywhere and also produced no calls in 300 families for each of 37 additional contexts. These comprised five evidence sources, ten network/walk variants, two architecture ablations and 20 disease relabellings. These diagnostics assess calling under the specified complete-null models.

Score-shift sensitivities extended the primary N0 experiment in **Methods, Assessment of calibration** and **Supplementary Fig. 8b,c**. Alternatives used N2 or the pair-weighted median of positive pilot standard deviations (SDs) within each weighted-degree pool, with isolated genes separate. Pools without positive SDs received no effective shift. N2 scales were re-estimated, so equal multipliers could represent different absolute increases. Nested candidate sets were shared across scales, strengths, methods and reference definitions. Both eligibility rules started from the original 7,763 candidates. Within each held-out configuration, the primary rule excluded focal-indication pseudo-seed genes; the sensitivity excluded genes seeded anywhere. Each rule used both unfiltered and focal-indication-seed-filtered references.

Comparators used empirical BH, empirical Benjamini–Yekutieli adjustment^4^, Gaussian-tail BH and a fixed raw-score cutoff of 0.0004493330539991 (**Supplementary Fig. 7a**; **Supplementary Table 5**). Adjusted-probability thresholds were 0.05. Gaussian upper-tail probabilities used each reference definition’s mean and sample SD. With zero SD, probabilities were zero above the constant reference value and one otherwise. Gaussian-tail testing differed from z-score ranking; the fixed raw-score cutoff had no false discovery rate (FDR) guarantee. The historical score-shift diagnostic, retained in **Supplementary Table 5 and Source Data,** added three reference-null SDs to a random 10% of eligible candidates in 300 held-out families. It used 4,000 other-block references and excluded genes seeded anywhere. Unlike current recall, historical recall included designated candidates with zero SD. Historical false discovery proportion (FDP) counted calls outside the designated set and was zero without calls. Mean recall was 0.7%; seven of 300 families produced calls.

Blockwise diagnostics applied BH *q* ≤ 0.05 separately to each 2,000-reference block and compared calls with the pooled 572-pair set (**Supplementary Fig. 8d**). Block agreement assessed reference-sampling sensitivity and was not required for primary support. The empirical-probability floor was $1/2,001$ per block and $1/6,001$ after pooling. Across 7,763 tests, the pooled floor implies a necessary minimum BH rejection rank of 26. Three of the original 20 disease-relabelled contexts had an empty block, illustrating finite-reference effects (**Supplementary Table 5**). Empirical support is assigned by randomized-reference comparisons and Benjamini–Hochberg adjustment under the specified testing choices. Empirical P values are reference-model tail probabilities; posterior false-pair probabilities require a separate probabilistic model. Candidate dependence and incomplete conditioning on seed topology constrain the calibration scope. FDR control when true signals are present, biological causality and sensitivity to biological effects require evidence beyond these diagnostics.

#### Supplementary Note 2. Temporal sensitivity analyses and event review

The clinical datasets were dated 22 December 2022 and 5 December 2025. STRING v12.0 was available in July 2023; repository history dated the disease-similarity artefact to October 2023, but the underlying MeSH release was undocumented. These dates establish pre-2024 input availability; the algorithm was not prospectively locked. The propagation engine and decision rule were developed after the outcome period began; 2023 served as a source-assembly interval. **Methods, Temporal evaluation of clinical outcomes** describes matching and exclusions. Match rates differed across support arms (**Supplementary Table 1**); the effect of differential matching on the temporal estimates remains unquantified.

**Sensitivity definitions and estimation.** Supplementary Fig. 6 defines any advancement and event-type-restricted advancement relative to the dated-advancement **endpoint in Methods**. In-window stage transitions were not verified for either sensitivity endpoint. The advancement amendment, introduced after arm-level feasibility tabulation to improve power, designated direct genetics as a positive control and BRIDGE-supported pairs as the headline comparison. It specified an inconclusive network-arm interpretation when its interval crossed one. The BRIDGE-supported comparison combined direct and network-added pairs against the common unprioritized group; network-added enrichment was assessed separately. Minimum detectable RS at 80% power was 3.1 for network-added launch and 1.7 for advancement. Calculation used a two-sided $\alpha=0.05$ Katz log-RS approximation with fixed arm sizes and comparator event rate.

Indication-cluster intervals used 5,000 resamples, retaining all pairs within sampled indications and taking percentiles of finite RS estimates, including zero. Indications containing network-added events were omitted in turn (**Supplementary Fig. 5**). Conditional logistic models used candidate pairs in the risk set, stratified by indication. Within-indication score percentiles were calculated from average ranks among candidate pairs in the risk set; higher percentiles denoted higher propagation scores. Models estimated odds ratios per 10-percentile-point increase with Wald intervals, then added $\log(1+\text{unweighted PPI degree})$ and categorical baseline phase. Of 5,807 eligible candidate pairs across 176 indications, 1,710 pairs across 25 indications with outcome variation contributed (**Supplementary** **Table 6**).

Exploratory activity and raw-score diagnostics were added after inspecting initial temporal results, without changing support assignments, eligibility or outcomes. Activity-stratification results retain estimates and contributing strata for the procedures in Methods. Sparse, differing subsets of contributing strata limit the assessment of activity-related confounding. These models estimate odds ratios. Raw-score selection fixed the top 572 of the original 7,763 candidates before temporal filtering, with T–I identifiers breaking ties. Filtering retained 345 network-added and 359 raw-score-selected pairs, sharing 227 pairs and all five network-added launches. Each rule used its own unselected remainder. Raw-score estimates and selection overlaps are provided. The two rules have overlapping selections and different comparators; their individual significance tests address separate contrasts.

**Review of network-added launch records.** The five pair entries mapped to three drug-programme records:

| **Network-added pair entries** | **Linked record** | **Recorded date and location** | **Interpretation** |
| --- | --- | --- | --- |
| *FGFR2*, *FGFR3*, *KDR*–nasopharyngeal neoplasms | Anlotinib/catequentinib | 28 April 2024, China | Three target pairs from one programme |
| *KIT*–gastrointestinal neoplasms | Imkeldi oral imatinib | 4 March 2025, USA | Formulation launch; event details concerned gastrointestinal stromal cancer, separately represented as historically launched |
| *SERPINC1*–haemophilia A | Fitusiran | 24 April 2025, USA | One target–indication entry |

The review supported annual window assignments for these five pair entries. Comprehensive validation of all 52 launches, non-events, matching and earliest-launch aggregation remains outstanding. January dates could be curated values rather than observed event days; cross-indication dates also occurred. The original workbook-to-curated reconstruction was unavailable. Further adjudication would require consistent rules across arms, including formulation identity and indication granularity. Event concentration and omission estimates appear in **Supplementary Fig. 5** and **Supplementary Table 6**.

#### Supplementary Figure 1

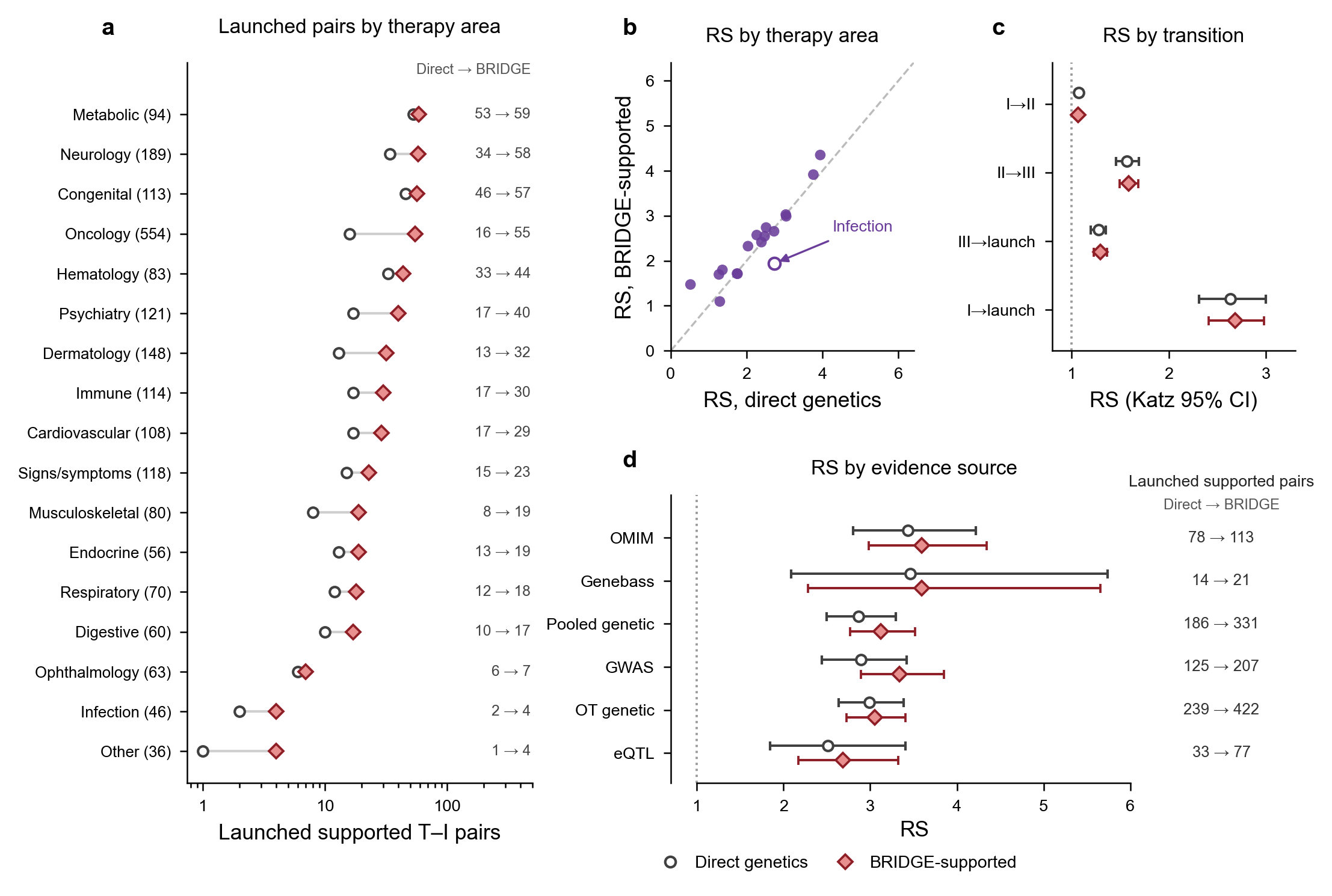

**Supplementary Figure 1 | Genetic support and clinical outcomes across therapy areas, development transitions and evidence sources.** BRIDGE-supported pairs combine directly supported and network-added pairs. **a,** Launched target–indication (T–I) pairs with direct genetic support (open circles) or BRIDGE support (red diamonds) across 17 Medical Subject Headings (MeSH) therapy areas. Grey lines connect counts within areas; adjacent values give supported counts and parentheses give all launched pairs. The x axis is logarithmic. Indications can belong to multiple areas, so counts are not additive. **b,** Phase-I-to-launch relative success (RS) for direct genetics and BRIDGE-supported pairs within each area. Purple points represent areas; the dashed diagonal marks equality. The open point denotes infection, with seven evaluable directly supported pairs. **c,** RS for phase-I-to-II, phase-II-to-III, phase-III-to-launch and phase-I-to-launch progression. **d,** Phase-I-to-launch RS across six separately calibrated genetic-evidence sources, each using its own seeded-indication context. OMIM, Online Mendelian Inheritance in Man; GWAS, genome-wide association study; OT, Open Targets; eQTL, expression quantitative trait locus. Adjacent counts give launched pairs supported by direct genetics and BRIDGE-supported pairs. Each RS compares a supported set with its complement in the corresponding evaluation population. In c,d, open circles and red diamonds denote direct genetics and BRIDGE-supported pairs, respectively. Horizontal lines show two-sided Katz 95% confidence intervals (CIs); dotted lines mark RS = 1.

#### Supplementary Figure 2

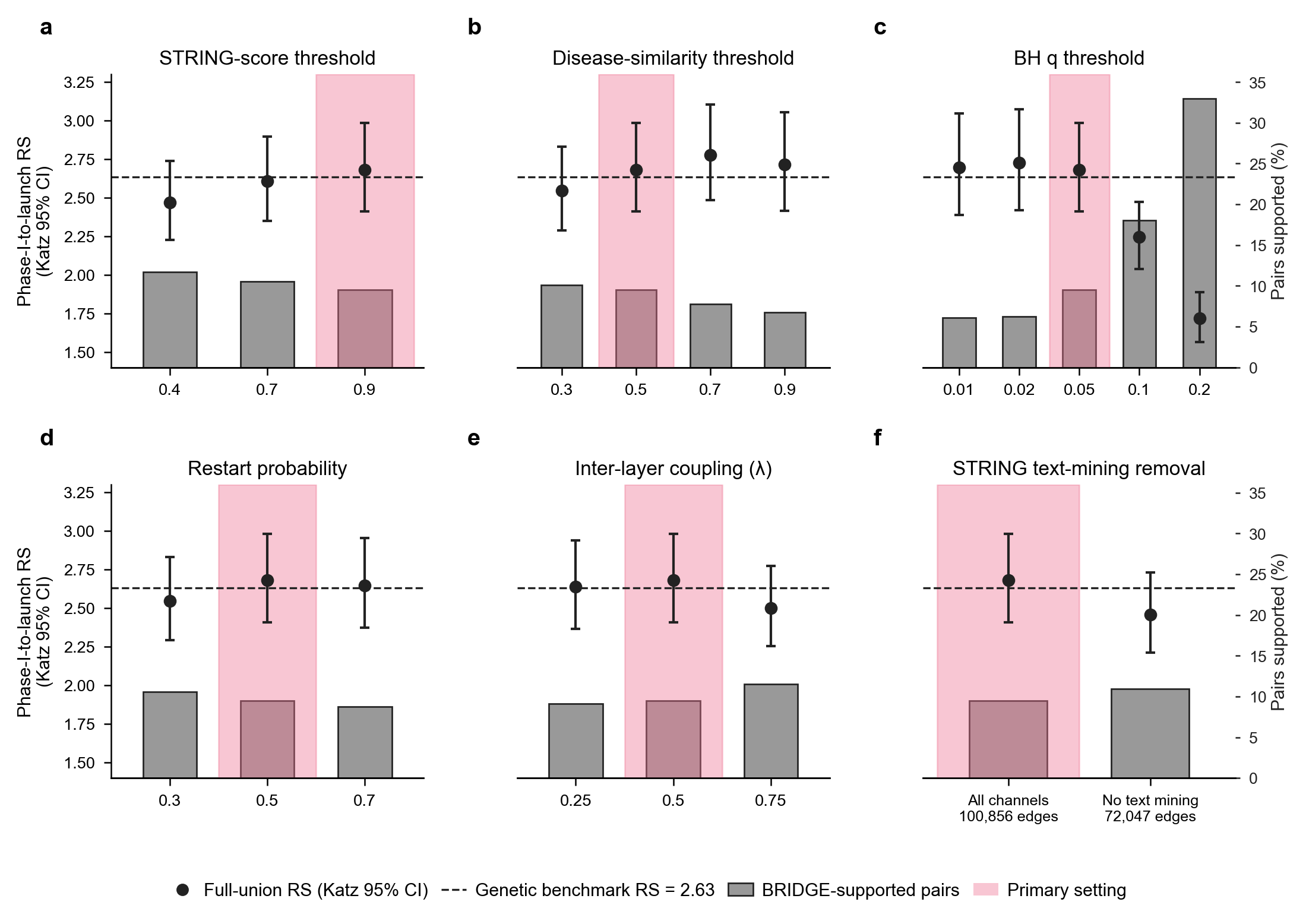

**Supplementary Figure 2 | Sensitivity of BRIDGE coverage and relative success to analysis settings.** BRIDGE-supported pairs combine directly supported and network-added pairs. Black circles and vertical lines show phase-I-to-launch relative success (RS) with two-sided Katz 95% confidence intervals (CIs), comparing each support set with its cohort complement. Grey bars show the percentage of the 13,022-pair cohort supported. The dashed line marks the direct-genetic RS of 2.63; pink shading identifies the primary setting. **a,** STRING combined-score threshold. **b,** Disease-similarity threshold. **c,** Benjamini–Hochberg (BH) *q* threshold. **d,** Restart probability. **e,** Inter-layer coupling parameter, λ. **f,** STRING networks with and without text-mining evidence, containing 100,856 and 72,047 retained edges, respectively. Each network or walk variant was recalibrated using its own 6,000-reference null bank. Panel c applies different *q* thresholds to the fixed primary BH-adjusted *q* values.

#### Supplementary Figure 3

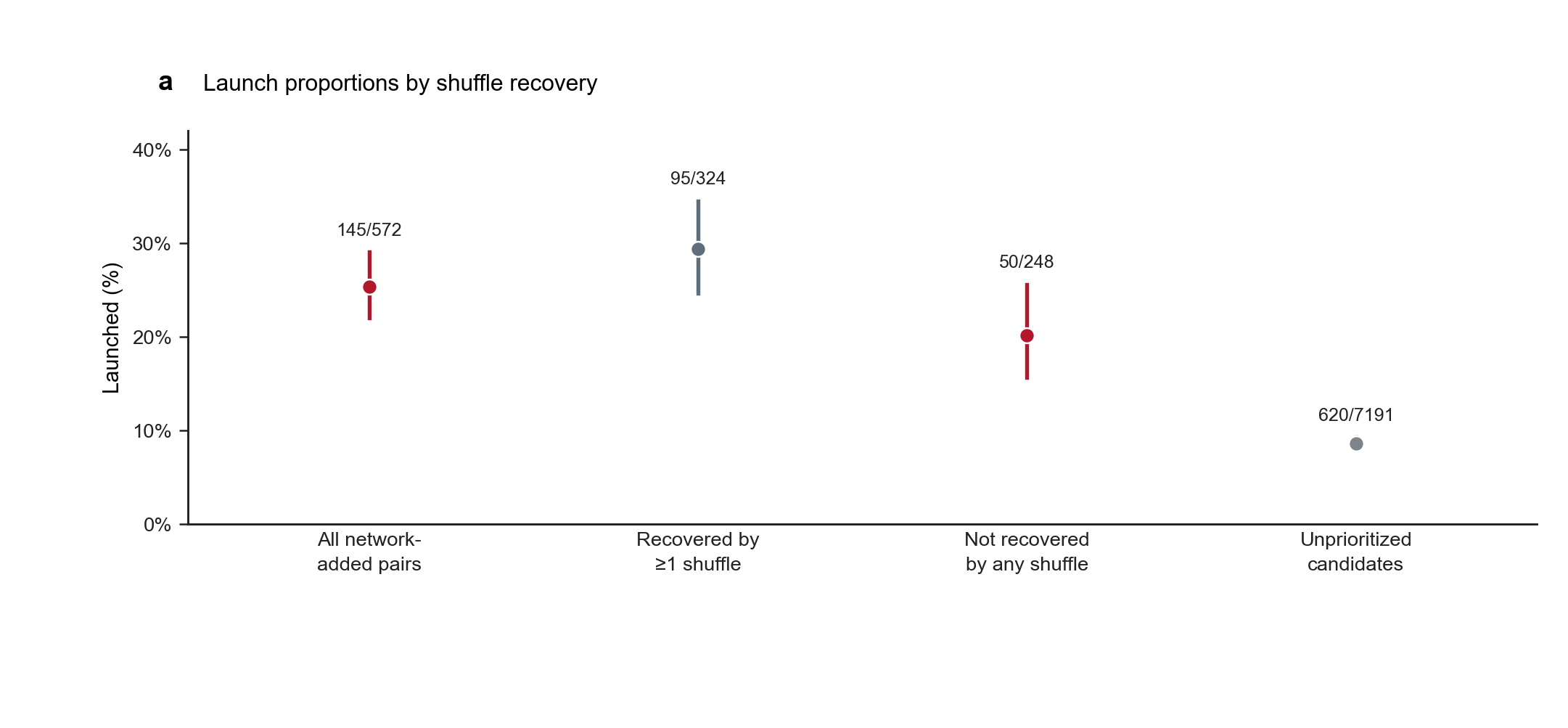

**Supplementary Figure 3 | Launch proportions by recovery under disease-label shuffling.** Observed and shuffled networks were separately recalibrated among the same 7,763 candidates, with support assigned at Benjamini–Hochberg q ≤ 0.05. Launch proportions among all 572 network-added pairs, those recovered by at least one of 100 shuffles (324 pairs), those never recovered (248 pairs), and unprioritized candidates (7,191 pairs). Points and lines show proportions and two-sided Wilson 95% confidence intervals; annotations give launched/total counts. Recovered and unrecovered groups partition the network-added pairs conditional on these 100 shuffles; the all-network-added group overlaps both. Across shuffles, the median fractions of each shuffle's supported pairs and unique implicated genes also present among the observed network-added pairs were 92.19% and 97.92%, respectively (denominators are the corresponding shuffled support sets). The original recovery histogram and overlap distribution are retained in Source Data under their original figure identifiers. Wilson intervals approximate independent pairs; shuffled-set overlaps are descriptive.

#### Supplementary Figure 4

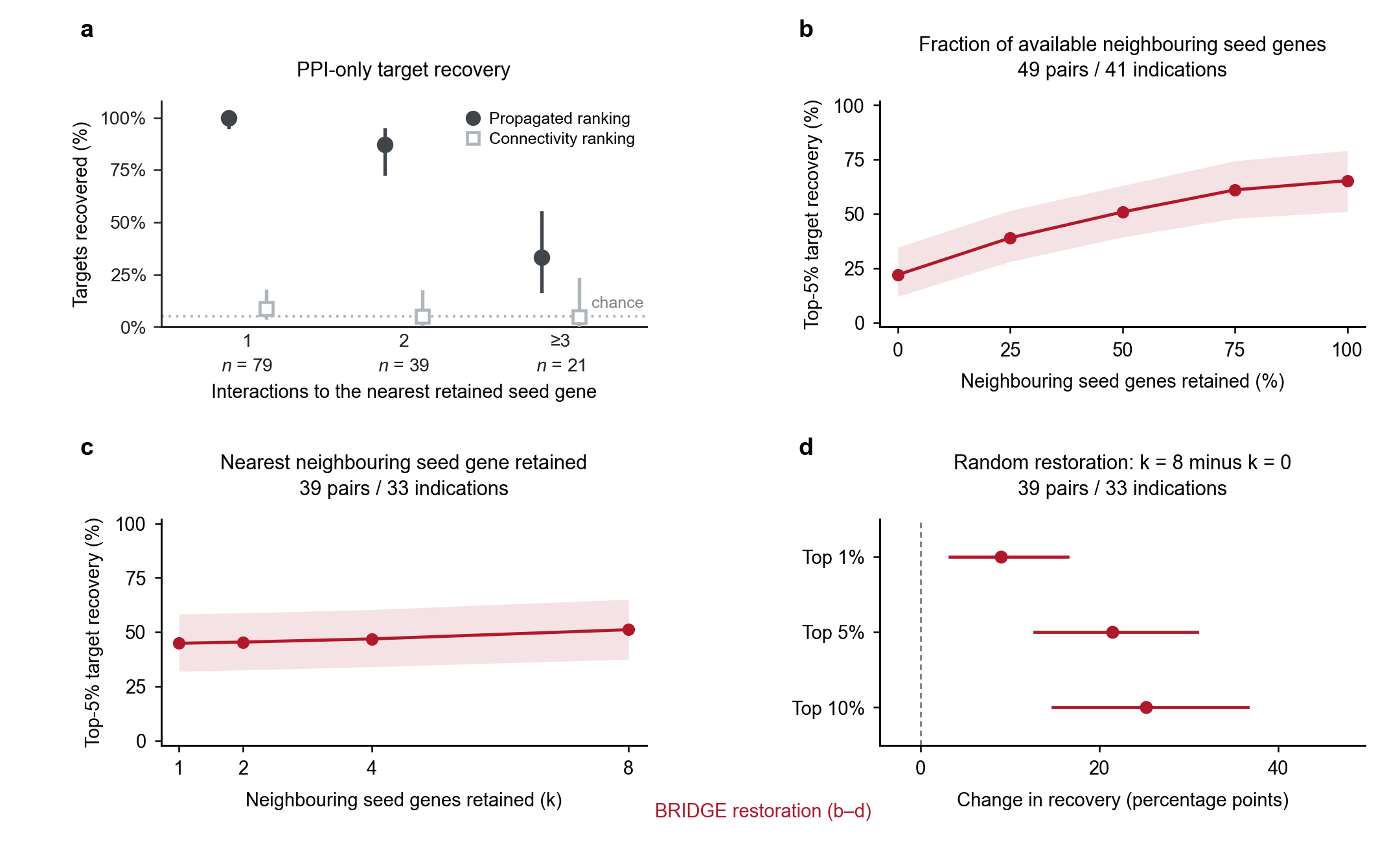

**Supplementary Figure 4 | Target withholding and sensitivity to restoring neighbouring seed genes.** **a,** Top-5% recovery after withholding each target’s seed assignment for the focal indication in 139 launched, directly supported pairs across 66 indications. Protein–protein interaction (PPI)-only propagation and unweighted PPI degree ranked the same genes with at least one retained PPI edge (non-isolated genes), excluding remaining seed genes. Points and vertical lines show recovery and two-sided Wilson 95% confidence intervals (CIs). Strata contain 79, 39 and 21 pairs at nearest-seed-gene distances 1, 2 and ≥3. The dotted line marks 5%; launch status determined eligibility only. **b–d,** BRIDGE recovery after withholding all focal-indication seed genes, for targets that are not seed genes elsewhere, and restoring neighbouring seed genes. Unrestored neighbouring seed genes received degree-stratum-matched surrogates under **Methods, Disease-context and target-recovery analyses**. **b,** Restoration of 0%, 25%, 50%, 75% or 100% of available neighbouring seed genes (49 pairs; 41 indications), rounding counts upwards. **c,** Nearest neighbouring seed gene retained at 1, 2, 4 or 8 restored (39 pairs; 33 indications with at least eight). Minimum distance to a neighbouring seed gene was fixed, but other distances and propagation mass could change. **d,** Paired changes from zero to eight randomly restored neighbouring seed genes at top-1%, top-5% and top-10% recovery in the same 39-pair cohort. Curves in b,c show top-5% recovery, averaging 50 matched backgrounds within pairs, then pairs. Bands in b,c and intervals in d show pointwise percentile CIs from 5,000 indication-cluster bootstrap resamples; d uses paired changes. All CIs are two-sided 95%. Targets in b–d were ranked among 19,699 genes. Zero restored neighbouring seed genes denotes surrogate replacement with other evidence retained. Recovery measures the fraction of eligible known-positive target–indication pairs whose targets rank within the specified top fraction.

#### Supplementary Figure 5

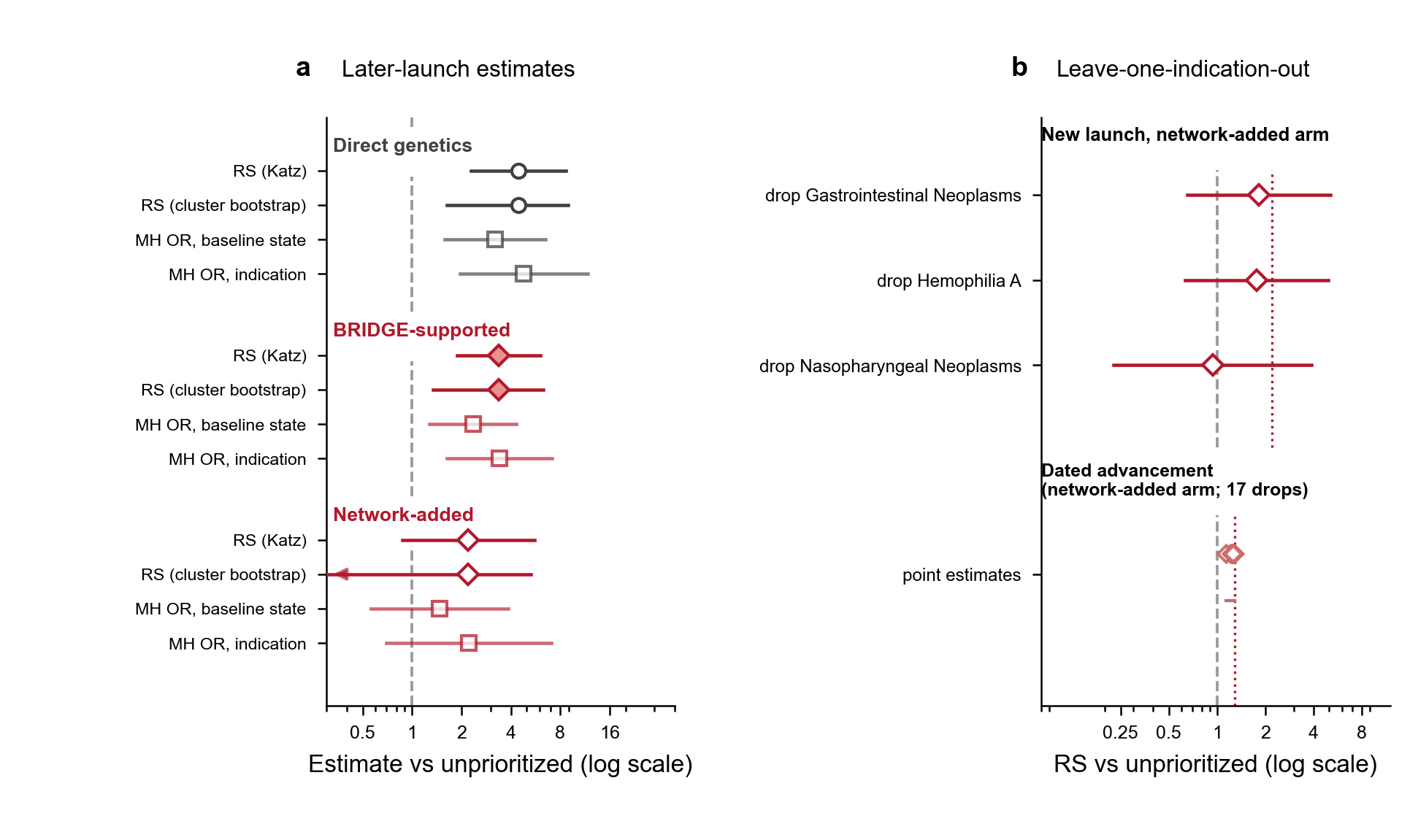

**Supplementary Figure 5 | Temporal outcome estimates and their concentration across indications.** The risk set contains 376 directly supported, 345 network-added and 5,462 unprioritized target–indication pairs; BRIDGE-supported pairs combine the first two groups. **a,** New-launch estimates against the common unprioritized group. Rows show unstratified relative success (RS; event-proportion ratio) with Katz intervals, RS with percentile intervals from 5,000 indication-cluster bootstrap resamples, and Mantel–Haenszel odds ratios (ORs; squares) stratified by baseline phase or indication. The arrow marks an interval extending to zero beyond the logarithmic axis. **b,** Network-added RS after omitting event-containing indications. Diamonds and horizontal lines show new-launch RS and Katz intervals for three indication omissions. For exploratory dated advancement, diamonds show 17 omission estimates and the short horizontal segment spans their range. Crimson dotted lines mark estimates before omission. Dated advancement required a higher recorded stage and a most recent event date in 2024–2025; that event need not identify the matched indication's stage transition. Grey dashed lines mark RS or OR = 1. All confidence intervals are two-sided 95% intervals. Bootstrap density histograms are retained in Source Data; the continuous-score models and contributing sample sizes are reported in Supplementary Table 6.

#### Supplementary Figure 6

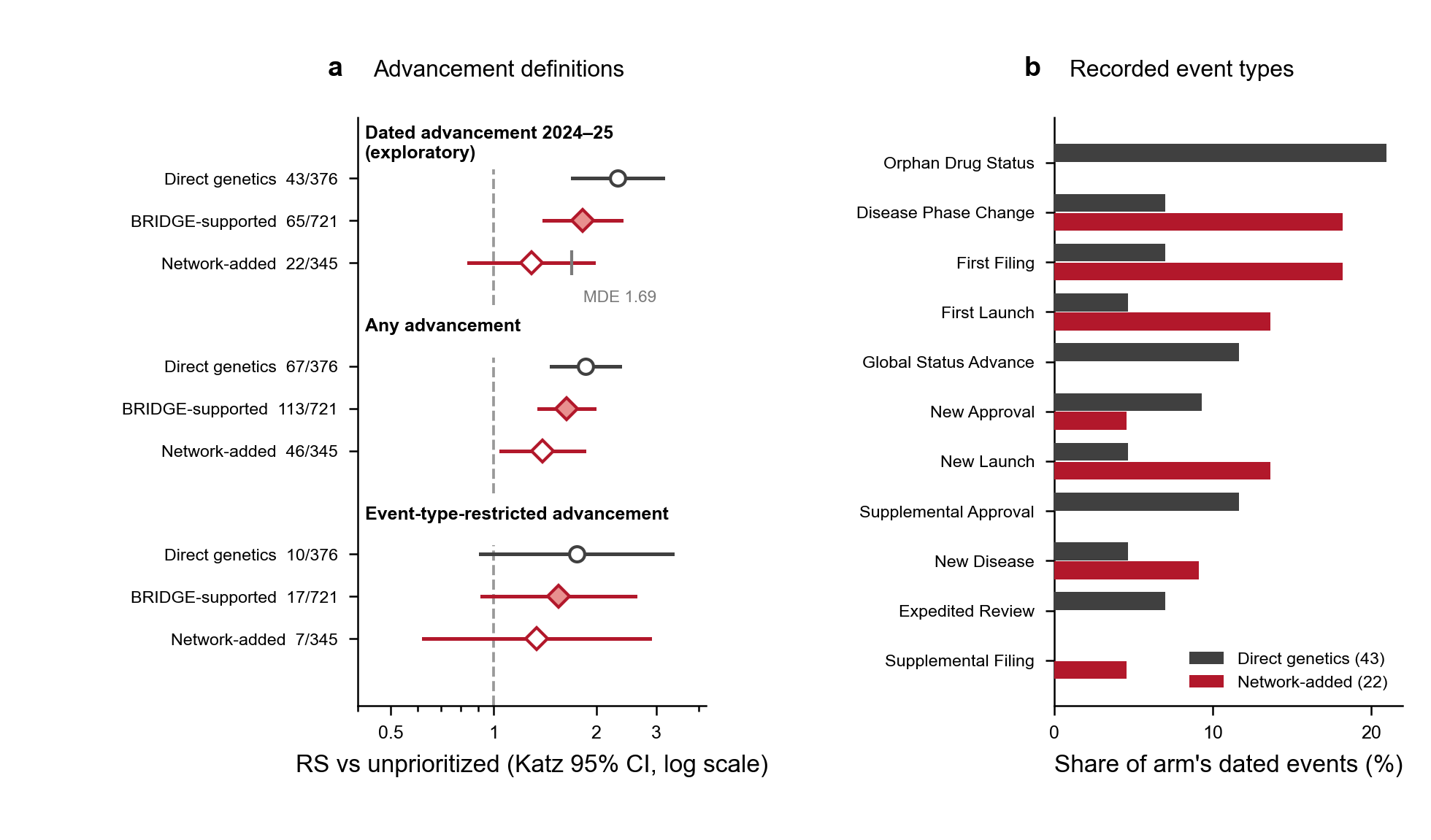

**Supplementary Figure 6 | Definitions and composition of clinical-advancement endpoints.** **a,** Relative success (RS; event-proportion ratio) for direct support, network-added support and BRIDGE-supported pairs against the common unprioritized group under three endpoint definitions. The risk set contains 376 direct, 345 network-added and 5,462 unprioritized pairs; BRIDGE-supported pairs combine the first two groups (721 pairs). Dated advancement required a higher recorded stage than the 22 December 2022 baseline and a most recent event date in 2024–2025. Any advancement omitted the date criterion. Event-type-restricted advancement combined dated advancement with Disease Phase Change, Global Status Advance or New Launch. The date could refer to another indication or an event without stage advancement. Points and horizontal lines show RS and two-sided Katz 95% confidence intervals; labels give events/pairs. The logarithmic axis has a dashed reference at RS = 1. The vertical tick marks the minimum detectable effect (MDE), expressed as RS, at 80% power for network-added dated advancement. **b,** Event types among dated-advancement pairs with direct support (43 pairs, black) or network-added support (22 pairs, red). Categories are the union of each group's six most frequent event types; percentages use each group's total dated-advancement pairs. Among pairs with any advancement, the numbers with a most recent event dated in 2024–2025 were 43/67 for direct genetics, 65/113 for BRIDGE-supported pairs, 22/46 for network-added pairs and 270/524 for unprioritized pairs. The original dated-share and indication-concentration displays are retained in Source Data.

#### Supplementary Figure 7

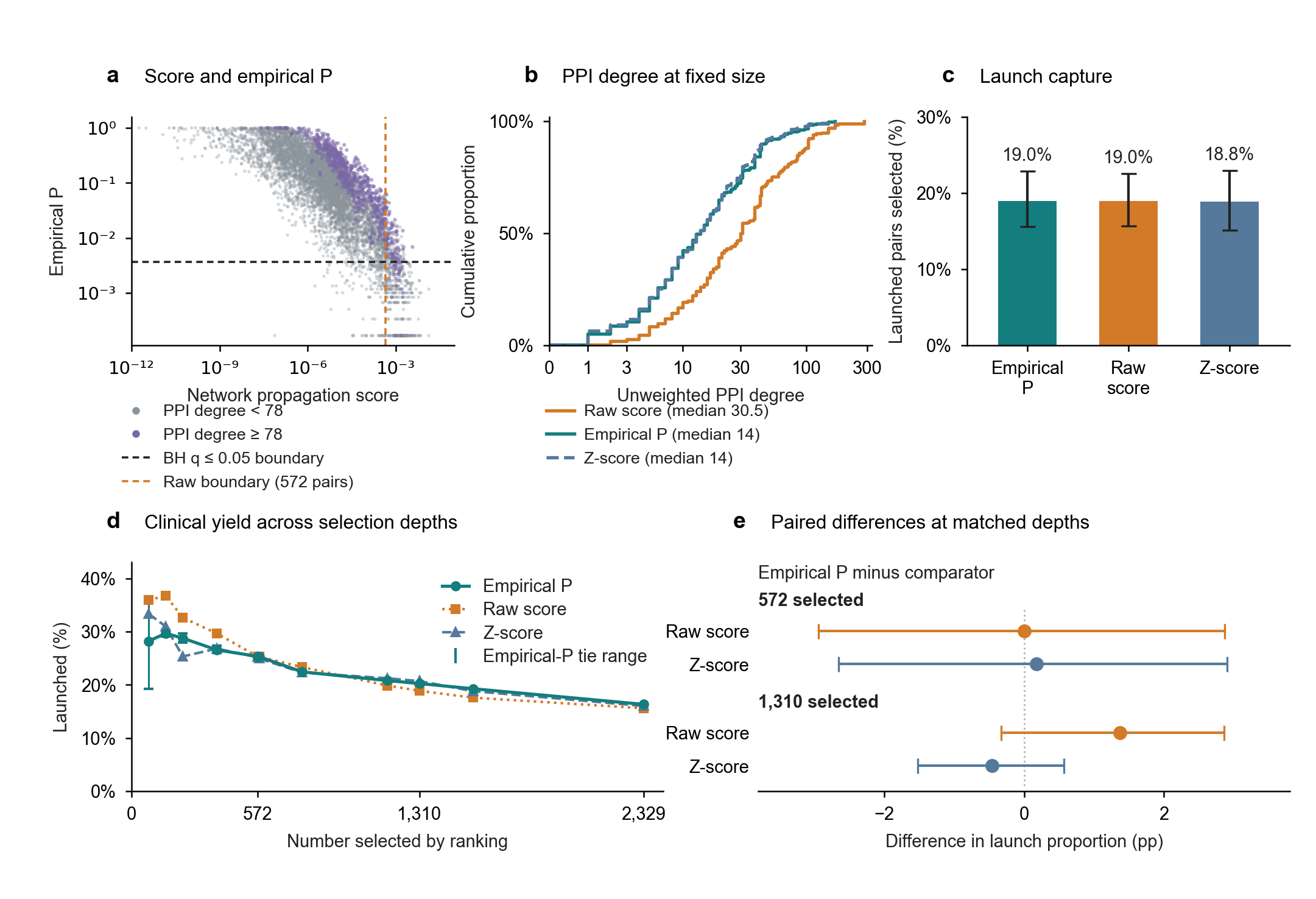

**Supplementary Figure 7 | Score normalization and clinical ranking comparisons.** **a,** Propagation scores versus candidate-specific empirical P values for 7,763 candidate target–indication pairs. Empirical P values used 6,000 degree-matched reference scores. Purple marks protein–protein interaction (PPI) degree ≥78 (90th percentile); grey marks lower degrees. Dashed boundaries mark the largest empirical P among 572 Benjamini–Hochberg (BH) q ≤ 0.05 network-added pairs (black; 0.003666) and the 572nd-highest raw score (orange; 0.0004493). Raw-score ties were resolved by pair identifier. Logarithmic axes omit 191 zero-score, uncalled pairs. **b,** Cumulative PPI-degree distributions among 572 pairs selected by empirical-P, raw-score and z-score rankings. Median degrees are 14, 30.5 and 14, respectively. The corresponding selections included 101, 90 and 113 distinct genes outside the 359-gene seed set. Degree + 1 is log-scaled; ticks show degree. **c,** Launch capture among all 765 launched candidates: 145/765, 145/765 and 144/765 for the respective rankings. Whiskers show two-sided pointwise 95% percentile confidence intervals from 5,000 shared bootstrap resamples of 205 indication clusters, conditional on fixed selections; each resample used its own launched-pair denominator. Empirical-P selection in b and c coincided with the network-added set; comparator selections were size-matched. **d,** Launch proportions across 11 selection depths. Teal whiskers show the full range across 200 alternative empirical-P tie resolutions, not sampling CIs. **e,** Empirical-P minus comparator launch proportions at 572 and 1,310 selected pairs. Points and whiskers show differences and pointwise, two-sided 95% CIs from 5,000 indication-cluster bootstrap resamples (205 clusters), conditional on fixed selections. Positive values favour empirical-P ranking; pp, percentage points. Selection at 1,310 is a ranking comparison.

#### Supplementary Figure 8

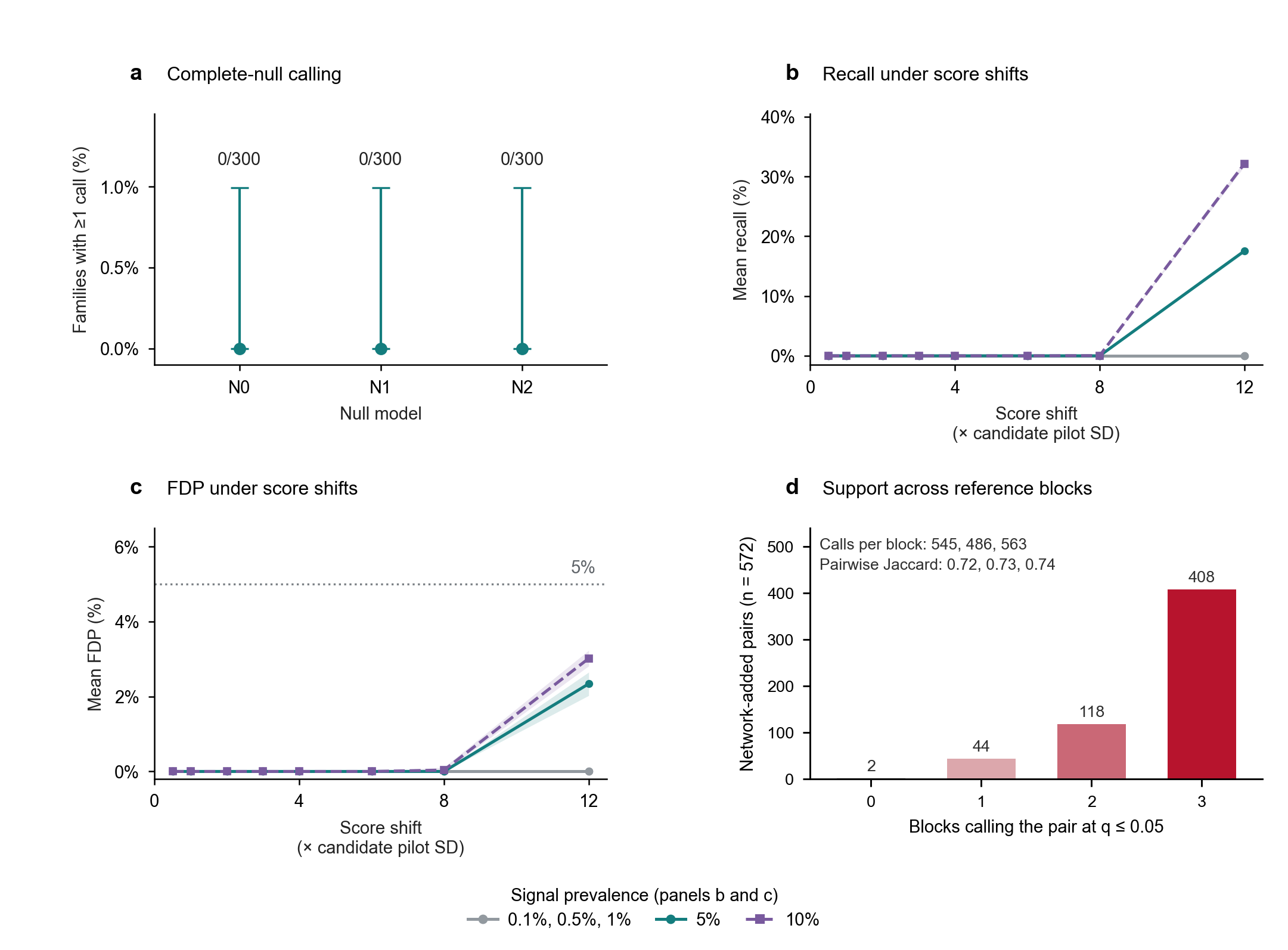

**Supplementary Figure 8 | Calibration diagnostics and reference-block stability.** **a,** Complete-null calling under degree-matched replacement (N0), a separate isolated-gene pool (N1), and replacement preserving seed-gene reuse across indications (N2). Each model used 6,000 references and 300 held-out families; candidates drawn as focal-indication seed genes in a held-out family were ineligible. Points show 0/300 families with any call; whiskers show exact one-sided 95% upper bounds. **b,** Mean recall under N0 across five signal prevalences and eight artificial score shifts (0.5–12 candidate-specific pilot standard deviations, SDs). Recall measures recovery of effective simulated signals (strict score increases); zero-shift candidates remain null and are excluded from recall denominators. **c,** Mean false discovery proportion (FDP), the fraction of calls without effective simulated signals, set to zero without calls. The dotted line marks 5%. In b and **c,** SDs came from 100 independent pilot families; each condition used 200 evaluation families, 6,000 references and BH q ≤ 0.05. Shading shows pointwise two-sided 95% percentile intervals from 5,000 family-bootstrap resamples, conditional on references and pilot scales. Sparse-prevalence curves (0.1%, 0.5%, 1%) coincide at zero; y scales differ. These panels summarize recovery and FDP under the specified artificial score shifts (Supplementary Note 1). **d,** Numbers of the 572 network-added pairs from the pooled 6,000-column reference bank also called by zero to three separate 2,000-column blockwise rules, each using q ≤ 0.05. Counts above bars partition the pooled set. Annotations show total calls per block and pairwise Jaccard indices in block-pair order 1–2, 1–3 and 2–3. Panel d shows complete observed counts; no confidence intervals are plotted for that panel.

#### Supplementary Figure 9

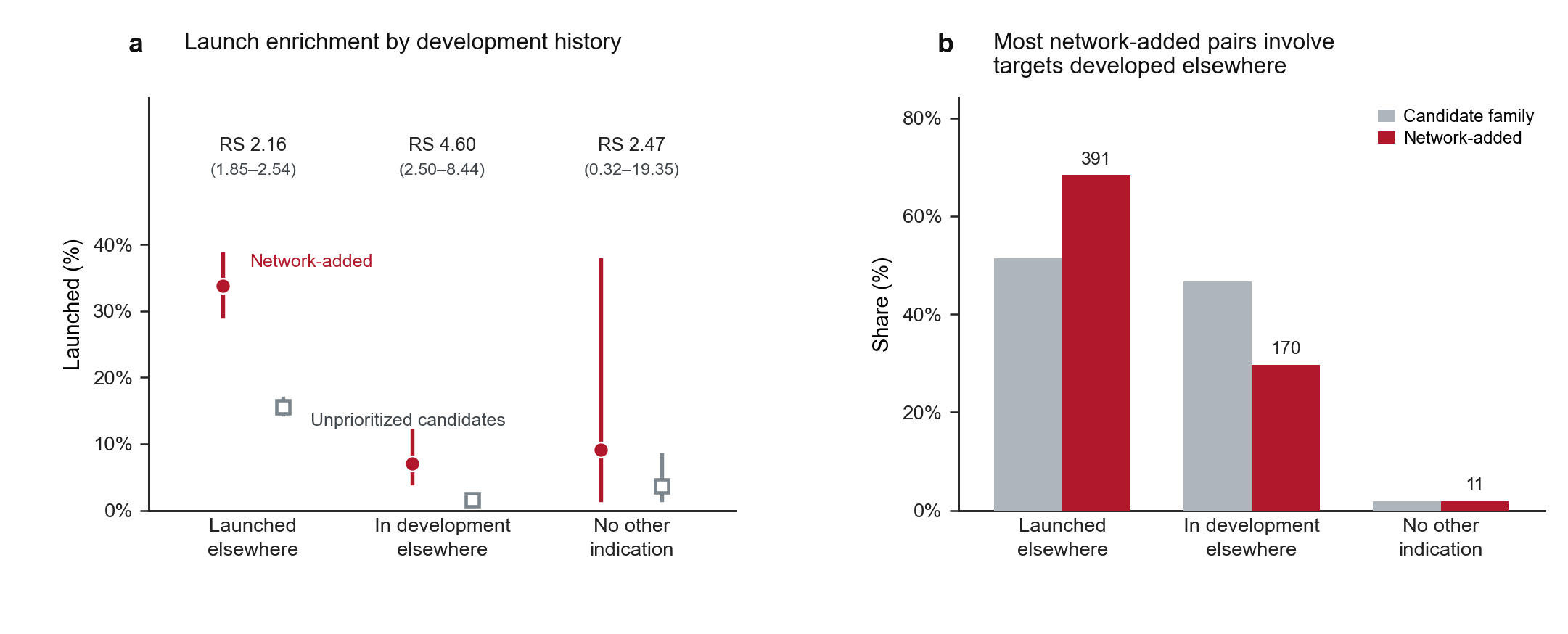

**Supplementary Figure 9 | Support and launch outcomes by development history in other indications.** Targets were grouped by their highest recorded phase outside the focal indication in the 13,022-pair benchmark: launched elsewhere, developed but unlaunched elsewhere, or represented only in the focal indication. **a,** Launch proportions for network-added pairs (red circles) and unprioritized candidates (open grey squares) within each group, with two-sided Wilson 95% confidence intervals (CIs). Annotations give relative success (RS), the supported/unprioritized launch-proportion ratio within each group, and two-sided Katz 95% CIs. **b,** Group proportions among 7,763 candidates (grey bars) and 572 network-added pairs (red bars); labels above red bars give network-added counts.

### Supplementary Table legends

**Supplementary Table 1 | Cohort and risk-set accounting.** Counts describe the retrospective target–indication cohort, mapped network context, clinical-status partition and temporal risk set. Count columns refer to pairs; proportions use the displayed denominators. Nine directly supported pairs outside STRING mapping remain in the full cohort but not the mapped network context. The mapped seed-gene and BRIDGE-supported gene sets contained 359 and 460 genes. Whole-cohort direct support contained 361 genes, including unmapped VDR and VEGFA, and the corresponding BRIDGE-supported gene set contained 462 genes.

**Supplementary Table 2 | Retrospective clinical enrichment and robustness.** Clinical transition counts and relative success (RS) for direct genetic support, BRIDGE-supported pairs, network-added support and robustness analyses. Each row identifies selected and reference groups; RS is the ratio of their event proportions. Confidence intervals are two-sided 95% using the stated method. Full curves, therapy-by-transition estimates and run-level details are in Source Data. Candidates whose genes were seeds for at least one other indication comprised 2,300 pairs (414 network-added; 1,886 unprioritized); candidates outside the seed-gene set comprised 5,463 pairs (158 network-added; 5,305 unprioritized). Each membership group used its own unprioritized remainder.

##### Same-target transfer comparison accompanying Supplementary Table 2

The four groups below partition all 7,763 candidates by same-target transfer eligibility and network-added support. Eligibility requires direct genetic support for the same target in a neighbouring indication. Within eligible pairs, RS was 1.75 (95% CI, 1.20–2.55). The workbook provides Wilson intervals for launch proportions and Katz intervals for RS; underlying records are in Source Data.

| **Same-target transfer eligibility** | **Network-added support** | **Launched/total pairs** | **Launch proportion** |
| --- | --- | --- | --- |
| Eligible | Supported | 74/290 | 25.5% |
| Eligible | Unprioritized | 32/220 | 14.5% |
| Ineligible | Supported | 71/282 | 25.2% |
| Ineligible | Unprioritized | 588/6,971 | 8.4% |

##### Ranking comparisons accompanying Supplementary Table 2

Among 175 candidates with zero reference variance, 174 had no observed excess and one ranked at positive infinity. Assigning all 175 the minimum finite z score changed outside-seed gene coverage slightly but retained 144 selected launches (**Supplementary Fig. 7**). Architecture comparisons covered all 11 selection depths. At 572 candidates, the full network had no established launch-proportion advantage over the alternatives; 13 of 100 disease-label shuffles matched or exceeded its launch count (*P* = 0.139). At 1,310 candidates, both architecture-difference intervals excluded zero and no shuffle reached the observed count (*P* = 1/101). At 1,310 pairs, all three score rankings also selected more launches than the local-transfer rule.

**Supplementary Table 3 | Network-added pairs outside the seed-gene set.** The 158 network-added pairs span 101 genes and include 38 launched pairs. Medical Subject Headings (MeSH) identifiers label indications. Pair-level empirical *P* values, Benjamini–Hochberg (BH)-adjusted *q* values and protein–protein interaction (PPI) topology are joined to gene-level annotations. Nearest focal-indication seed genes and representative full-network routes identify available connections; they do not quantify contributions to propagation scores. Blank distances and “No focal-indication PPI path” denote no path in that relation. Support without text mining denotes support after STRING text-mining exclusion and recalibration (81 of 158 pairs). Tractability flags indicate source-resource membership, as detailed below.

##### Catalogue annotations and case evidence accompanying Supplementary Table 3

Before final trait–indication matching, 85 of the 101 added genes satisfied the source-specific genetic criteria described in **Methods**, excluding somatic evidence. These source-level annotations precede indication-specific matching.

Antibody and small-molecule flags record membership in the gene lists released by Minikel et al.^5^, including medium-to-low-confidence antibody predictions; flag absence denotes absence from those lists. The suitability of each therapeutic modality (such as an antibody or small molecule), efficacy and safety require indication-specific assessment. Catalogue launch status refers to supported pairs in the historical dataset. Development records describe the recorded history across indications; current activity and undrugged opportunities require separate assessment.

| **Supported pair** | **Genetic-source and network detail** | **Intervention evidence and interpretation** |
| --- | --- | --- |
| *ALAS1*–porphyrias (D011164); *q* = 0.03329 | OMIM *HMBS*–acute intermittent porphyria (D017118) matched porphyrias at similarity 0.864; HMBS–ALAS1 edge weight, 0.918. | RNAi evidence concerns acute intermittent porphyria within the broader catalogue indication.^6,7^ ALAS1 lacked both catalogue modality flags. |
| *TFPI*–haemophilia A (D006467) and B (D002836); each *q* = 0.00995 | OMIM *F10*–factor X deficiency (D005171) matched A/B at 0.843/0.809. PICCOLO factor VII traits also supported *F10* for A; the *F7* seed gene used factor VII traits (D005168) at 0.817. Exact OMIM *F8*–A and *F9*–B seed genes were two steps away through F10. TFPI–F10, F10–F8 and F10–F9 weights were 0.999, 0.997 and 0.953. | Matched F10/F7 seed genes were not haemophilia-specific causal associations. TFPI inhibition provides compensatory intervention.^8^ BASIS compared marstacimab with participants' preceding treatment periods in haemophilia A/B without inhibitors, rather than randomized placebo treatment.^9^ |
| *IL23A*–rheumatoid arthritis (D001172); *q* = 0.03349 | Supported pair retained as a negative therapeutic example. | Guselkumab did not improve the primary rheumatoid-arthritis endpoint in a randomized phase 2 trial.^10^ |

The ALAS1 and both TFPI pairs retained support after text-mining exclusion. Three network-added *HCAR2* pairs lacked a focal-indication PPI path, although all 158 catalogue pairs had a full-network route to genetic evidence. STRING edges encode functional associations, including indirect relationships. Case-source records and seed-gene routes accompany the catalogue. Cases were selected retrospectively, with scores, clinical outcomes and temporal endpoints fixed.

##### Catalogue candidates beyond the local-transfer rule

Joining the 158-pair catalogue to the frozen local-transfer ledger identified 129 pairs involving 87 genes with a direct PPI connection to a focal-indication seed gene. The remaining 29 pairs involved 20 genes and satisfied neither branch of the rule. Six genes contributed pairs to both groups. Beyond-rule distances were two PPI steps for 16 pairs, at least three for 10 and no focal-indication PPI path for three. Text-mining exclusion retained eight of 29 beyond-rule pairs and 73 of 129 local-rule pairs. These summaries describe model sensitivity. The complete catalogue, 29 beyond-rule pairs and sensitivity settings accompany Source Data.

##### IFNLR1–COVID-19 source and therapeutic evidence

*IFNLR1*–COVID-19 (D000086382) was supported in the primary analysis (*q* = 0.0334917) and after text-mining exclusion (*q* = 0.0408285). Its sole focal-indication seed gene, *IFNAR2*, was two PPI steps away. Four qualifying Open Targets Genetics records matched COVID-19 exactly (trait–indication similarity 1): GCST010936, GCST011073, GCST90095123 and GCST90095124, with normalized locus-to-gene shares of 0.62, 0.55, 0.61 and 0.58, respectively. The records include overlapping consortium analyses and are not four independent replications. GCST011073 is a release 5 dataset linked by the GWAS Catalog to a 2020 consortium article; that publication year is not the release date. Original qualifying records and case routes provide provenance. The representative IFNLR1–IFNA1–IFNAR2 STRING path traces functional associations rather than a receptor-binding or signalling sequence. The pair remained supported without disease-similarity edges but not under PPI-only propagation.

Santer et al.^11^ examined samples from a prior randomized phase 2 study. Single-cell profiles from nine participants (five treated, four placebo) showed interferon-stimulated gene responses in IFNLR1-expressing peripheral immune cells following pegylated interferon lambda. This secondary sample analysis supports the IFNLR1–IL10RB receptor-pathway interpretation. Reis et al.^12^ tested a single 180-μg dose within seven days of symptom onset in predominantly vaccinated adults. The primary composite of COVID-19 hospitalization or emergency observation exceeding six hours within 28 days occurred in 25/931 versus 57/1,018 participants (relative risk, 0.49; 95% Bayesian credible interval, 0.30–0.76). In an earlier 120-participant trial, Jagannathan et al.^13^ found no shortening of viral shedding (median seven days in both groups; adjusted hazard ratio, 0.81; 95% CI, 0.56–1.19) or symptom duration. The studies evaluated different endpoints and settings, with therapeutic evidence concerning the IFNLR1–IL10RB receptor complex. The frozen 22 December 2022 catalogue already recorded active phase III development for this pair, and the phase 3 publication preceded its selection for this manuscript.

##### Exploratory sensitivity to seed-gene replacement

The primary protocol was fixed before scoring all 29 beyond-rule pairs across 26 indications. Seed-gene replacement procedures are specified in **Methods, Target catalogue and genetic anchors**. Reconstruction and convergence checks are retained in Source Data, seed-replacement numerical checks. Gene-rank percentiles compare each target with all 19,699 mapped genes; lower percentiles indicate better ranks. Median replacement scores were below the originals for all 29 pairs. Median rank worsened for 24 pairs and improved for five. Because replacement also changes other genes’ scores, lower target scores can accompany better ranks. The median of pair-specific median score ratios was 0.8405; the median rank-percentile change was +0.1015 percentage points. Score and rank effects were heterogeneous and generally smaller than in the IFNLR1 example. Pair summaries, individual configurations and surrogate identities are retained.

For *IFNLR1*, replacing the sole *IFNAR2* seed gene shifted its rank percentile from 0.12945% to a median of 22.4935% (5th–95th perturbation percentiles, 2.5565–35.8648%) and yielded a median replacement/original score ratio of 1.9637 × 10^–5^. All 100 controls were three to six PPI steps away, so this comparison also changed the distance to the nearest seed gene. A separate exploratory amendment, fixed after that diagnostic but before further scoring, evaluated all 15 eligible controls in the same degree stratum and exactly two PPI steps from *IFNLR1*. Their median rank percentile was 0.70816% (full range, 0.18021–1.75897%) and median score ratio was 0.11507 (full range, 0.03901–0.34193). All 15 scores were positive and lower than the original, while all ranks remained in the top 1.8%. The complete distance-matched controls are supplied. Matching distance substantially attenuated the rank shift; the remaining differences may reflect precise degree, edge weights, alternative paths and other network features. Reported ranges summarize computational perturbations; *P* values, *q* values and support assignments were not recalculated.

**Supplementary Table 4 | Recovery after withholding and restoring seed genes.** Recovery in the top 5% under target withholding, indication withholding and seed-gene restoration. Pair counts and Wilson 95% confidence intervals summarize withholding. Restoration values are means across Monte Carlo backgrounds with saved indication-cluster bootstrap intervals; the stratum label distinguishes count-scale and fraction-scale doses.

**Supplementary Table 5 | Calibration and null-model diagnostics.** Diagnostic estimates under the original degree-matched null (N0), separate isolated-gene-pool null (N1) and one-to-one replacement null preserving seed-gene reuse across indications (N2), together with null-block replication and historical planted-score diagnostics. Complete-null bounds use the source method specified. Selected score-shift conditions summarize sparse-signal and strongest-shift behaviour. Complete condition grids, historical diagnostics and per-family records are retained in Source Data.

**Supplementary Table 6 | Temporal clinical-outcome estimates.** New launch is the primary outcome; advancement endpoints are exploratory. Counts, relative success (RS), model odds ratios and two-sided 95% intervals are presented by support contrast and estimation method. The continuous-score model is scaled per 10 within-indication score-percentile points, with higher percentiles indicating higher propagation scores. Detailed event definitions, leave-one-indication-out estimates and indication-concentration tables are retained in Source Data.
